# Assessment of the Utility of Gene Positioning Biomarkers in the Stratification of Prostate Cancers

**DOI:** 10.1101/728972

**Authors:** Karen J. Meaburn, Tom Misteli

## Abstract

In eukaryotic cells, the genome is spatially organized in a non-random fashion within the confines of the interphase nucleus, and most genes occupy preferred nuclear positions. For some genomic loci these positioning patterns are context specific, reflected in the distinct location of certain genes and chromosomes in different cell types and in disease. Disease-related differential spatial positioning of genes has led to the hypothesis that the spatial reorganization of the genome may be utilized as a diagnostic biomarker. In keeping with this possibility, the positioning patterns of specific genes can be used to reproducibly discriminate benign tissues from cancerous ones. In addition to the use of spatial genome organization for diagnostic purposes, we explore here the potential use of spatial genome organization as a prognostic tool. This is a pressing need since in many cancer types there is a lack of accurate markers to predict the aggressiveness of individual tumors. We find that directional repositioning of *SP100* and *TGFB3* gene loci stratifies prostate cancers of differing Gleason scores. A more peripheral position of *SP100* and *TGFB3* in the nucleus, compared to benign tissues, is associated with low Gleason score cancers, whereas more internal positioning correlates with higher Gleason scores. Conversely, *LMNA* is more internally positioned in many non-metastatic prostate cancers, while its position is indistinguishable from benign tissue in metastatic cancer. Our findings of subtype-specific gene positioning patterns in prostate cancer provides a proof-of-concept for the potential usefulness of spatial gene positioning as a prognostic biomarker.

## Introduction

The genome is highly spatially organized within the interphase nucleus (Cremer and Cremer, 2001;Bickmore, 2013). Most chromosomes, genes, and individual non-coding regions of the genome occupy preferred nuclear positions relative to the center of the nucleus or to other nuclear landmarks, such as associations with other genomic loci or nuclear bodies (Takizawa et al., 2008b;Bickmore and van Steensel, 2013;Meaburn, 2016). Some loci alter their position under different physiological conditions, for example, between cell/tissue types (Boyle et al., 2001;Parada et al., 2004;Peric-Hupkes et al., 2010;Meaburn et al., 2016) or between different proliferation states (Bridger et al., 2000;Meaburn and Misteli, 2008;Chandra et al., 2015). Spatial reorganization of the genome is also a common feature of disease, and has been documented in a wide range of pathologies, including epilepsy (Borden and Manuelidis, 1988), Down syndrome (Paz et al., 2015), laminopathies (Meaburn et al., 2007;Mewborn et al., 2010), viral and parasitic infections (Li et al., 2010;Knight et al., 2011), and cancer (Meaburn, 2016;Mai, 2018). Repositioning events are loci-specific and do not reflect global genome reorganization events (Meaburn, 2016).

Although the spatial organization of the genome has been studied for decades, how gene positioning patterns are established and maintained remains largely elusive. It is also unclear if the nuclear position of a locus is important for function or is largely a consequence of nuclear activities (Meaburn, 2016). Most often, a functional link is drawn between spatial genome organization and gene expression (Brown et al., 1997;Brickner and Walter, 2004;Williams et al., 2006;Takizawa et al., 2008a;Peric-Hupkes et al., 2010), however, there are also many instances where changes in gene expression and nuclear position of a locus are unrelated (Scheuermann et al., 2004;Williams et al., 2006;Kumaran and Spector, 2008;Meaburn and Misteli, 2008;Harewood et al., 2010;Meaburn, 2016). Most likely, there are multiple mechanisms in play to determine the spatial organization of the genome (Shachar et al., 2015;Meaburn, 2016;Randise-Hinchliff et al., 2016). In addition to gene expression, chromatin modifications, even in the absence of changes in gene expression (Towbin et al., 2012;Therizols et al., 2014;Harr et al., 2016;Cabianca et al., 2019;Falk et al., 2019), replication timing (Hiratani et al., 2008) and a variety of structural nuclear proteins (Dundr et al., 2007;Meaburn et al., 2007;Solovei et al., 2013;Zuleger et al., 2013;Shachar et al., 2015) have be implicated in the positioning of genomic loci.

While the mechanisms governing spatial positioning patterns are unclear, the fact that the genome is spatially reorganized in disease begs the question of whether spatial positioning patterns can be exploited for clinical purposes (Meaburn, 2016;Mai, 2018). We have previously demonstrated that the positioning patterns of some genes can be used to reproducibly and accurately discriminate benign breast and prostate tissues from cancerous ones (Meaburn et al., 2009;Leshner et al., 2016;Meaburn et al., 2016). For instance, the positioning patterns of *HES5* and *FLI1* are highly indicative of cancer, with both *HES5* and *FLI1* repositioned in 100% of breast cancers and *FLI1* repositioned in 92.9% of prostate cancers, compared to benign tissue controls (Meaburn et al., 2009;Leshner et al., 2016;Meaburn et al., 2016). High repositioning rates result in low false negative detection rates. Crucially for diagnostic applications, many of the genes that reproducibly reposition in cancer show limited variability between morphologically normal tissues and do not reposition in benign disease, yielding low false positive detection rates (Meaburn et al., 2009;Leshner et al., 2016;Meaburn et al., 2016). Given the sensitivity and specificity for the positioning patterns of several genes in detecting cancer, these small-scale studies suggest gene positioning biomarkers (GPBs) could be a useful addition to cancer diagnostics.

Prostate cancer is a leading cause of cancer and cancer-related deaths (Bray et al., 2018). As with most cancers, while there is value in additional diagnostic biomarkers, there is also a critical need for additional prognostic biomarkers, to predict best treatment options, including to reduce overtreatment in patients whose cancer would have remained asymptomatic during their lifetime without treatment (Welch and Black, 2010;Sandhu and Andriole, 2012). Currently, the cornerstone of predicating a patient’s outcome is the Gleason grading system, which is based on histological assessment (Epstein et al., 2005;Brimo et al., 2013). In this system the architectural structure of the prostate tissue is graded from Gleason grade 1, which represents a well differentiated tissue morphology, to the very poorly differentiated Gleason grade 5. The two most prominent Gleason grades in a given tumor/biopsy are summed to give a Gleason score (Epstein et al., 2005;Epstein, 2018). Low Gleason score cancers are more likely to be indolent, whereas higher Gleason scores correlates with poor outcomes (Albertsen et al., 1998;Pound et al., 1999;Brimo et al., 2013). However, further markers are required as there is a range in outcomes for patients with the same Gleason score.

To improve on the Gleason system, additional clinical factors, most commonly serum prostate-specific antigen (PSA) levels, T stage (size of tumor/spread to nearby tissues), percentage of cancer positive biopsy cores, and patient age are taken into account (D’Amico et al., 1998;Thompson et al., 2007;Cooperberg et al., 2009;Chang et al., 2014). ~15% of patients are diagnosed with high-risk (likely to cause morbidity, recur, metastasize and/or be lethal) prostate cancer, based on PSA levels of >20ng/ml and/or Gleason score of 8-10 and/or T stage of either T2c-T4 or T3a-T4, depending on the classification system (Thompson et al., 2007;Cooperberg et al., 2009;Chang et al., 2014). Low-risk cancers (PSA <10ng/ml, Gleason score 2-6, and T stage T1-T2a) are generally predicted to remain asymptomatic during the patient’s lifetime and the use of active surveillance/watchful waiting is often recommended, as opposed to active treatment (Thompson et al., 2007;Cooperberg et al., 2009). Conversely, intermediate risk (Gleason score 7 or T stage T2b/c) patients generally receive treatment (Thompson et al., 2007;Cooperberg et al., 2010). Gleason scores can be subject to inter- and intra-observer variability, usually of just a single Gleason score (Montironi et al., 2005), but for patients at the border of low and intermediate risk this may make the difference of receiving treatment or not. Moreover, with the current clinical criteria to stratify risk, both over- and under-treatment remains a concern for all prostate cancer risk groups (Cooperberg et al., 2010;Punnen and Cooperberg, 2013). Improved markers are needed to better distinguish indolent from high-risk prostate cancers and to aid classification of intermediate-risk cancers, to reduce overtreatment and optimize therapeutic strategies.

There is a growing number of genomic prognostic biomarkers for prostate cancer, including several commercial assays based on the DNA methylation status of a small number of genes or on gene expression (Kornberg et al., 2018). Additionally, changes to nuclear size and shape and gross chromatin texture, which are not considered in Gleason scoring, provide additional predictive power to detect aggressive prostate cancers (Veltri and Christudass, 2014;Hveem et al., 2016). Few studies have assessed the prognostic potential of the spatial organization of the genome. The most compelling evidence for prognostic GPBs comes from telomeres, where increased telomeric aggregation correlates with progression and risk in several types of cancers (Mai, 2018). Similarly, in a single acute myeloid leukemia patient, HSA8 and 21 became more proximal to each other while the patient was in remission, prior to a disease relapse and re-emergence of t(8;21) in the patient’s bone marrow (Tian et al., 2015).

Here, we explore the utility of spatial gene positioning patterns to identify clinically distinct subgroups of prostate cancer. We find subtype-specific positioning for *SP100*, *TGFB3* and *LMNA*. The direction in which *SP100* and *TGFB3* reposition, compared to benign tissue, distinguishes low and intermediate/high Gleason score cancers, whereas *LMNA* repositions in many non-metastatic cancers but not in metastatic cancers. Although the sensitivity of this assay is currently too low to be clinically useful, our findings of subtype-specific gene positioning patterns in prostate cancer provides additional evidence for the potential of spatial genome organization as a novel prognostic biomarker.

## Materials and Methods

### Tissue Fluorescence *in situ* Hybridization (FISH)

4-5μm thick normal, benign disease (hyperplasia and chronic prostatitis), and cancerous formalin-fixed, paraffin embedded (FFPE) human prostate tissues were obtained from US Biomax Inc, Imgenex Corporation, BioChain Institute, or the University of Washington (Prof. Lawrence True) under the guidelines and approval of the Institutional Review Board of the University of Washington (#00-3449) (Supplementary Table 1). Patient tissues were de-identified before receipt.

To generate probe DNA for FISH, bacterial artificial chromosome (BAC) clones were labeled with biotin- (Roche), digoxigenin- (Roche) or DY-547P1- (Dyomics GmbH) conjugated dUTPs by nick translation (Meaburn, 2010). The following BACs were used: RP11-727M18 (to position *SP100*, chromosome location: 2q37.1); RP11-270M14 (*TGFB3*, 14q24.3); RP11-1021J5 (*SATB1*, 3p24.3); RP11-35P22 (*LMNA*, 1q22) (BACPAC resource center). Single- or dual-probe FISH experiments were performed as previously described (Meaburn et al., 2009;Leshner et al., 2016;Meaburn et al., 2016), with the following modifications: for most tissues a 1 hour 60ºC bake step was performed, prior to the xylenes (Macron Fine Chemicals) deparaffinization step; 40μg yeast RNA (Life Technologies) was used in place of tRNA; DyLight 488 labeled anti-digoxigenin (Vector Laboratories) was occasionally used to detect digoxigenin-labeled probe DNA; and no probe detection steps were required for the fluorescently labeled DY-547P1-dUTP FISH probes.

### Image Acquisition

Epithelial nuclei were randomly imaged throughout the tissue, unless benign and malignant glands were present in the same tissue section. In such cases, care was taken to image and analyze the different morphologies separately, whilst still acquiring epithelial cell nuclei randomly within the benign or malignant regions to capture as much diversity within the cancer (or benign tissue) as possible. Image accusation was performed as previously described, using an IX70 (Olympus) Deltavision (Applied Precision) system, with a 60x 1.42N oil objective lens (Olympus), an auxiliary magnification of 1.6, and a X-Y pixel size of 67.25nm (Meaburn et al., 2009;Leshner et al., 2016;Meaburn et al., 2016) or with a similar imaging regime using an IX71 (Olympus) Deltavision (Applied Precision) system, 100x 1.40N oil objective lens (Olympus), with an X-Y pixel size of 64.6nm. Image stacks were acquired to cover the thickness of the tissue section, with a 0.5μm or 0.25μm step interval along the Z axis, respectively. All image stacks were deconvolved and converted to maximum intensity projections using SoftWoRx (Applied Precision). The change in acquisition approach did not affect the resulting positioning data from the image datasets. We obtained similarly statistically identical distributions for the position of a gene in a given tissue using the two different acquisition methods (*P* = 0.79-0.86, Kolmogorov-Smirnov (KS) test), as from repeat analysis of tissues using an identical acquisition method (*P* = 0.65-0.99 (Meaburn et al., 2009), unpublished data).

### Image Analysis

Image analysis to determine the radial position of a gene within a tissue was performed as previously described (Meaburn et al., 2009;Leshner et al., 2016;Meaburn et al., 2016). Briefly, 96-167 interphase epithelial nuclei were manually segmented in Photoshop (Adobe) for each gene in each tissue, except for *TGFB3* in tissue C10 where 88 nuclei were segmented. To map the radial position of the gene loci, nuclei were run though custom image analysis software scripts, using MATLAB (The Mathsworks Inc.), with DIPImage and PRTools toolboxes (Deft University, P. Gudla and S Lockett (NCI/NIH); (Meaburn et al., 2009)). Euclidean distance transform (EDT) was computed for each nucleus, to assign every pixel within the nucleus its distance to the nearest nuclear boundary. The software then determined the nuclear EDT position of the geometric gravity center of the automatically detected FISH signals. To normalize for variations in nuclear size and shape between specimens, the EDT of a FISH signal was normalized to the maximal nuclear EDT for that nucleus, with 0 denoting the nuclear periphery and 1 the nuclear center. The normalized FISH signal EDTs for a given gene in each specimen was then combined to produce a relative radial distribution (RRD), and a cumulative frequency distribution was generated. All detected alleles in a nucleus were included, regardless of the number present. In the case of the pooled normal distributions (PNDs), the normalized FISH EDTs from all allele in all the normal tissues analyzed, for a given gene, were combined into a single dataset (Supplementary Figure 1). The number of nuclei and tissues used in each PND were as follows: the *SP100* PND contained 845 nuclei from 7 normal tissues; *TGFB3*, 996 nuclei from 8 normal tissues; *SATB1*, 874 nuclei from 7 normal tissues; and *LMNA*, 725 nuclei from 6 normal tissues. Finally, to statistically compare a gene’s positioning patterns, RRDs between tissues, or between specimens and the PND, were cross-compared using the nonparametric two-sample 1D KS test, where *P* < 0.01 was considered significant.

Some previously reported RRDs were included in the current analysis (Supplementary Table 1; (Leshner et al., 2016;Meaburn et al., 2016)), which were compared to an update PND. The four PNDs used in this study included normal tissues N6 and N7, in addition to the normal tissues previously reported, which did not affect the RRDs (*P* = 0.83-1, 1D KS test). RRDs for *TGFB3* in tissues C25, C27, B9, N3, N4, N11-14 were previously reported in (Meaburn et al., 2016), and the RRDs of *SP100* in C11, C12, C13, C25, N1, N2, N6-10, *SATB1* in C11, C12, C13, C25, B9, N1, N2, N6-10, and *LMNA* in C11, C18, C19, N5, N10 and N15-16 were reported in (Leshner et al., 2016).

## Results

### Mapping of Candidate Genes in Prostate Tissues

We have previously identified genes that radially reposition in breast and/or prostate cancer and have demonstrated their potential as diagnostic biomarkers (Meaburn et al., 2009;Leshner et al., 2016;Meaburn et al., 2016). Here, we sought to extend these studies to determine if candidate genes occupied distinct nuclear positions between different subgroups of prostate cancer, with the goal of assessing their utility for cancer prognostics. To identify prognostic candidate genes we took advantage of our previous studies, in which we had screened the radial positions of 47 genes in a panel of prostate cancers (Leshner et al., 2016;Meaburn et al., 2016). From that gene set we chose two genes, *SATB1* and *LMNA*, for further assessment as potential biomarkers of high-risk prostate cancer because both genes repositioned in a single high-risk T3 stage cancer, but not in two intermediate risk T2 cancers, or a low risk T2 cancer (Leshner et al., 2016). We also selected *SP100* to test its potential as a marker of low risk, since we previously found it to reposition in a low risk Gleason score 6 prostate cancer, but not in three intermediate or high-risk Gleason score 7 cancers (Leshner et al., 2016). Finally, we selected *TGFB3* for further analysis since it repositioned in one of two low risk Gleason score 6 prostate cancers, but not in two prostate cancers of unknown Gleason score and TNM stage (Meaburn et al., 2016), representing a potential low-risk/indolent prostate cancer biomarker.

To determine whether the positioning patterns of these genes were able to stratify prostate cancers into clinically relevant subgroups, we performed FISH on a panel of 4-5μm thick FFPE human prostate tissues, which included a diverse group of 32 prostate cancer specimens covering a range of Gleason scores and T stages, with and without known metastases, and 25 benign prostate tissues (for details see Supplementary Table 1). To map the spatial positioning pattern of a gene in a given tissue, we measured the radial position, normalized for nuclear size and shape, of each locus in ~120 epithelial interphase nuclei as previously described (see Materials and Methods; (Meaburn et al., 2009;Leshner et al., 2016;Meaburn et al., 2016). The normalized radial position of each gene was determined and the cumulative RRDs were statistically compared to a PND, a standardized normal distribution created by pooling all nuclei from normal tissues for a given gene, or individual tissues using the 1D KS test, with *P* < 0.01 considered significant (see Materials and Methods, (Meaburn et al., 2009;Leshner et al., 2016;Meaburn et al., 2016), Supplementary Figure 1).

We initially assessed the repositioning rates for the candidate genes in the assorted set of prostate cancer samples. Compared to the PND, *SP100* was in a statistically significantly different radial position in 44.4% (12/27) prostate cancer specimens (Figure 1, Table 1, Supplementary Table 2). Similarly, *LMNA* repositioned in 36.4% (4/11), *SATB1* in 34.8% (8/23), and *TGFB3* in 31.8% (7/22) of prostate cancer tissues (Figure 1, Table 1, Supplementary Tables 3-5). The repositioning rates are slightly higher than in the previous smaller scale studies (25-33.3%) (Leshner et al., 2016;Meaburn et al., 2016). However, in keeping with previous findings, all four genes repositioned in too few cancers to be of use as prostate cancer diagnostic GPBs since detecting cancer based on the repositioning of the gene would misclassify 55.6-68.2% of the tumors as not cancerous, depending on the gene. The likelihood of a gene repositioning in a cancer did not correlated with gene copy number (Figure 1, Supplementary Table 6).

**Figure 1.**
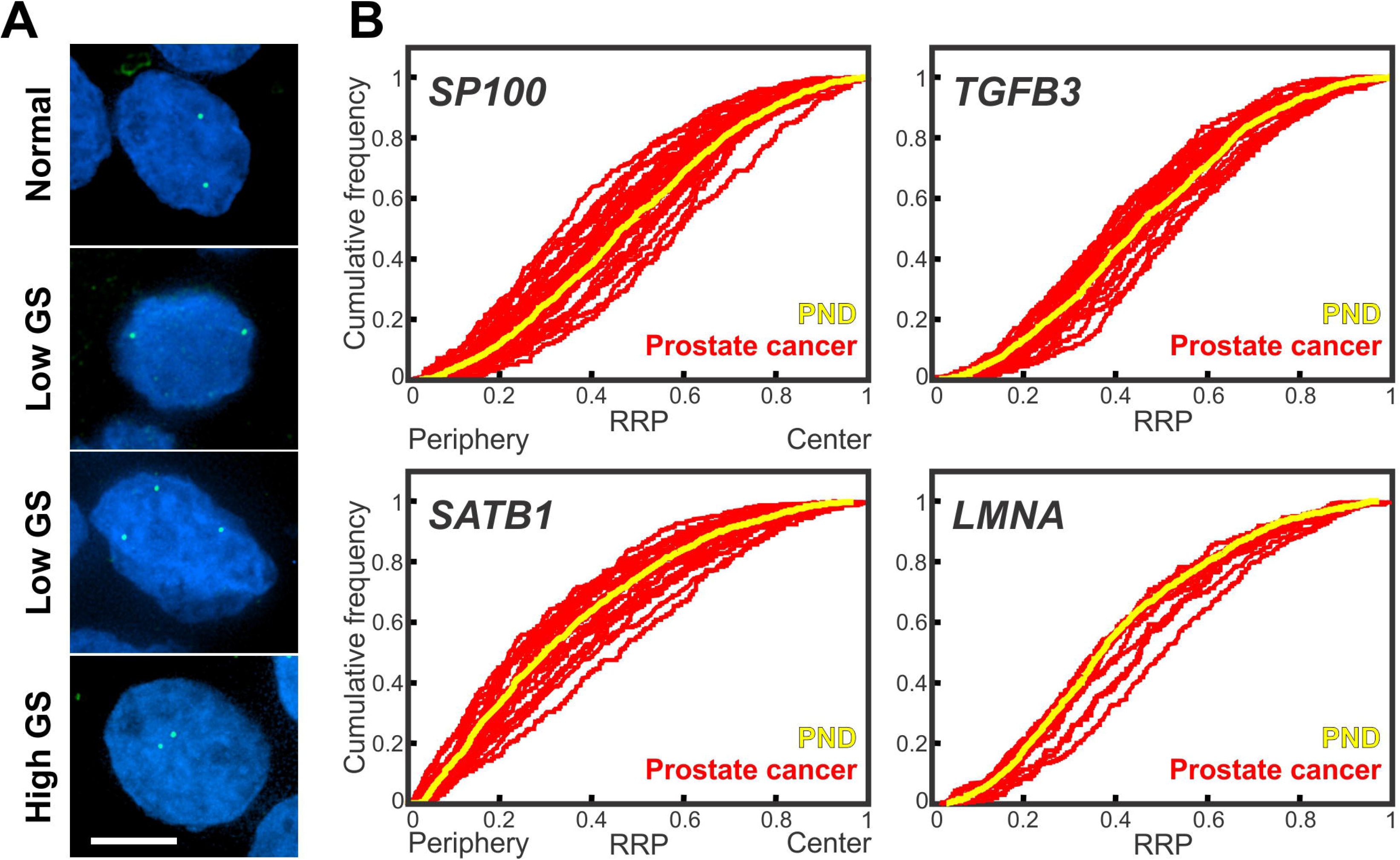
Gene positioning in prostate cancer tissues. **(A)** Gene loci were detected by FISH in FFPE prostate tissue sections. *SP100* gene loci (green) in normal and cancerous prostate tissues. GS, Gleason score. Projected image stacks shown. Nuclei were counterstain with DAPI (blue). Scale bar, 5μm. **(B)** Cumulative RRDs for the indicated genes in prostate cancer (red) and the pooled normal distribution (PND; Yellow). RRP, relative radial position.

**Table 1.**
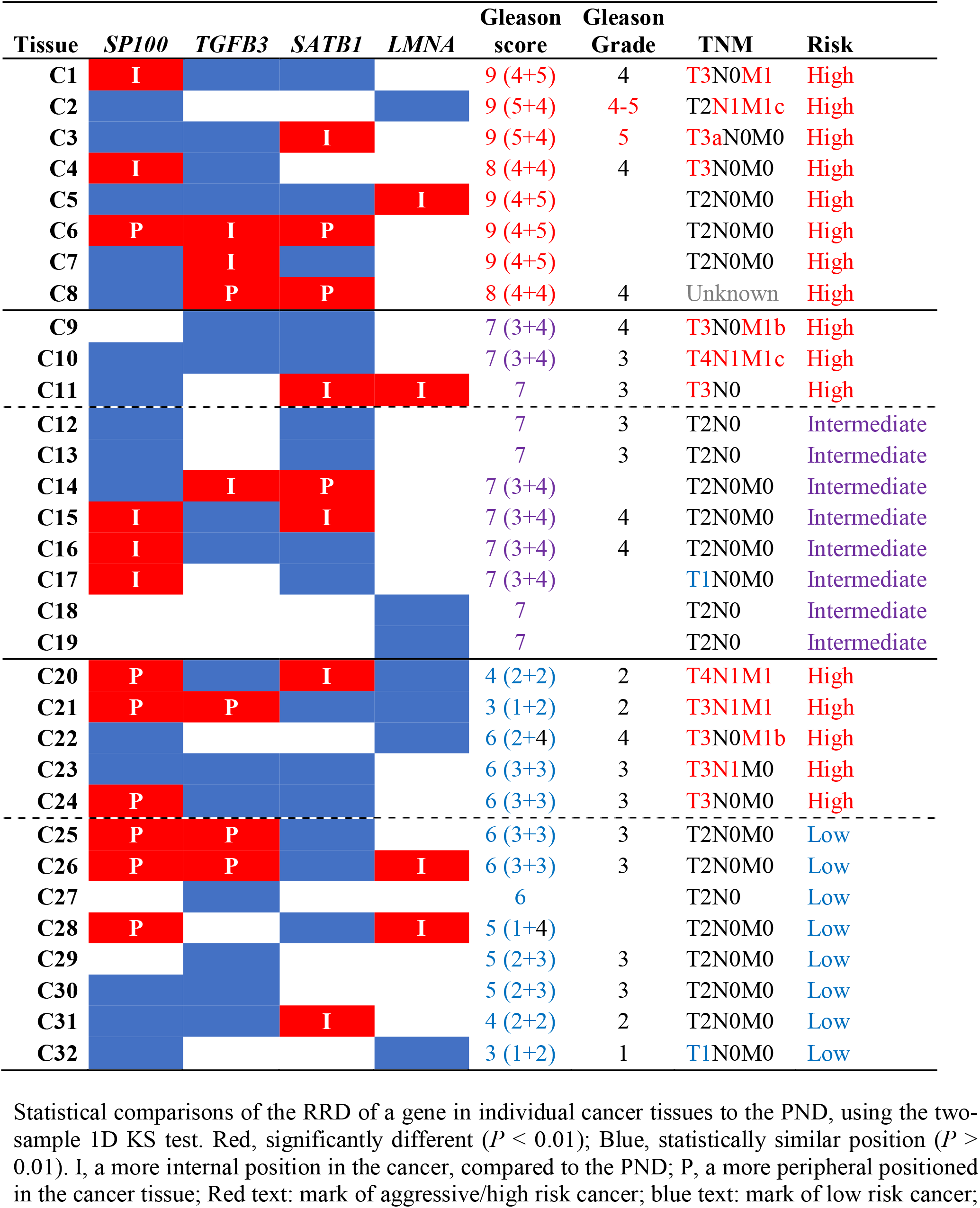
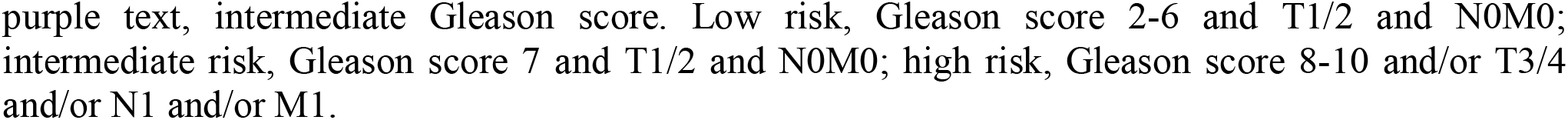
Spatial repositioning of target genes in prostate cancer.

| Tissue | *SP100* | *TGFB3* | *SATB1* | *LMNA* | Gleason score | Gleason Grade | TNM | Risk |
| --- | --- | --- | --- | --- | --- | --- | --- | --- |
| C1 | I |  |  |  | 9 (4+5) | 4 | T3N0M1 | High |
| C2 |  |  |  |  | 9 (5+4) | 4-5 | T2N1M1c | High |
| C3 |  |  | I |  | 9 (5+4) | 5 | T3aN0M0 | High |
| C4 | I |  |  |  | 8 (4+4) | 4 | T3N0M0 | High |
| C5 |  |  |  | I | 9 (4+5) |  | T2N0M0 | High |
| C6 | P | I | P |  | 9 (4+5) |  | T2N0M0 | High |
| C7 |  | I |  |  | 9 (4+5) |  | T2N0M0 | High |
| C8 |  | P | P |  | 8 (4+4) | 4 | Unknown | High |
| C9 |  |  |  |  | 7 (3+4) | 4 | T3N0M1b | High |
| C10 |  |  |  |  | 7 (3+4) | 3 | T4N1M1c | High |
| C11 |  |  | I | I | 7 | 3 | T3N0 | High |
| C12 |  |  |  |  | 7 | 3 | T2N0 | Intermediate |
| C13 |  |  |  |  | 7 | 3 | T2N0 | Intermediate |
| C14 |  | I | P |  | 7 (3+4) |  | T2N0M0 | Intermediate |
| C15 | I |  | I |  | 7 (3+4) | 4 | T2N0M0 | Intermediate |
| C16 | I |  |  |  | 7 (3+4) | 4 | T2N0M0 | Intermediate |
| C17 | I |  |  |  | 7 (3+4) |  | T1N0M0 | Intermediate |
| C18 |  |  |  |  | 7 |  | T2N0 | Intermediate |
| C19 |  |  |  |  | 7 |  | T2N0 | Intermediate |
| C20 | P |  | I |  | 4 (2+2) | 2 | T4N1M1 | High |
| C21 | P | P |  |  | 3 (1+2) | 2 | T3N1M1 | High |
| C22 |  |  |  |  | 6 (2+4) | 4 | T3N0M1b | High |
| C23 |  |  |  |  | 6 (3+3) | 3 | T3N1M0 | High |
| C24 | P |  |  |  | 6 (3+3) | 3 | T3N0M0 | High |
| C25 | P | P |  |  | 6 (3+3) | 3 | T2N0M0 | Low |
| C26 | P | P |  | I | 6 (3+3) | 3 | T2N0M0 | Low |
| C27 |  |  |  |  | 6 |  | T2N0 | Low |
| C28 | P |  |  | I | 5 (1+4) |  | T2N0M0 | Low |
| C29 |  |  |  |  | 5 (2+3) | 3 | T2N0M0 | Low |
| C30 |  |  |  |  | 5 (2+3) | 3 | T2N0M0 | Low |
| C31 |  |  | I |  | 4 (2+2) | 2 | T2N0M0 | Low |
| C32 |  |  |  |  | 3 (1+2) | 1 | T1N0M0 | Low |

Statistical comparisons of the RRD of a gene in individual cancer tissues to the PND, using the two-sample 1D KS test. Red, significantly different ( $P < 0.01$ ); Blue, statistically similar position ( $P > 0.01$ ). I, a more internal position in the cancer, compared to the PND; P, a more peripheral positioned in the cancer tissue; Red text: mark of aggressive/high risk cancer; blue text: mark of low risk cancer;
purple text, intermediate Gleason score. Low risk, Gleason score 2-6 and T1/2 and N0M0; intermediate risk, Gleason score 7 and T1/2 and N0M0; high risk, Gleason score 8-10 and/or T3/4 and/or N1 and/or M1.

In addition to whether a gene was repositioned, the direction of its repositioning was also determined (Figure 1, Table 1). Of the 12 prostate cancers in which *SP100* was repositioned, the gene was more internally positioned compare to the PND in five cancer tissues (5/12; 41.7%) and more peripherally positioned in seven (58.3%). Similarly, *TGFB3* was more internally positioned in three (3/7; 42.9%) cancers and more peripherally positioned in four (57.1%). *SATB1* was more internally positioned in five of the eight cancers where the gene was repositioned (62.5%) and more peripherally positioned in three cancers (37.5%). Conversely, *LMNA* repositioned to a more internal nuclear location in all four cancer specimens in which the gene was repositioned (Figure 1, Table 1). The direction of repositioning accounted for most of the differences in the positioning patterns for a given gene between the cancer tissues in which repositioning occurred. There was little variation between the RRDs of a gene between the cancers in which the gene was more internally positioned. Similarly, there was little statistical variation in RRDs among cancers in which the gene was more peripherally positioned, with the exception of *SATB1* (Table 2, Supplementary Tables 2-5). Taken together, we find heterogeneity in the radial positioning patterns for all four candidate genes between prostate cancers.

**Table 2.**
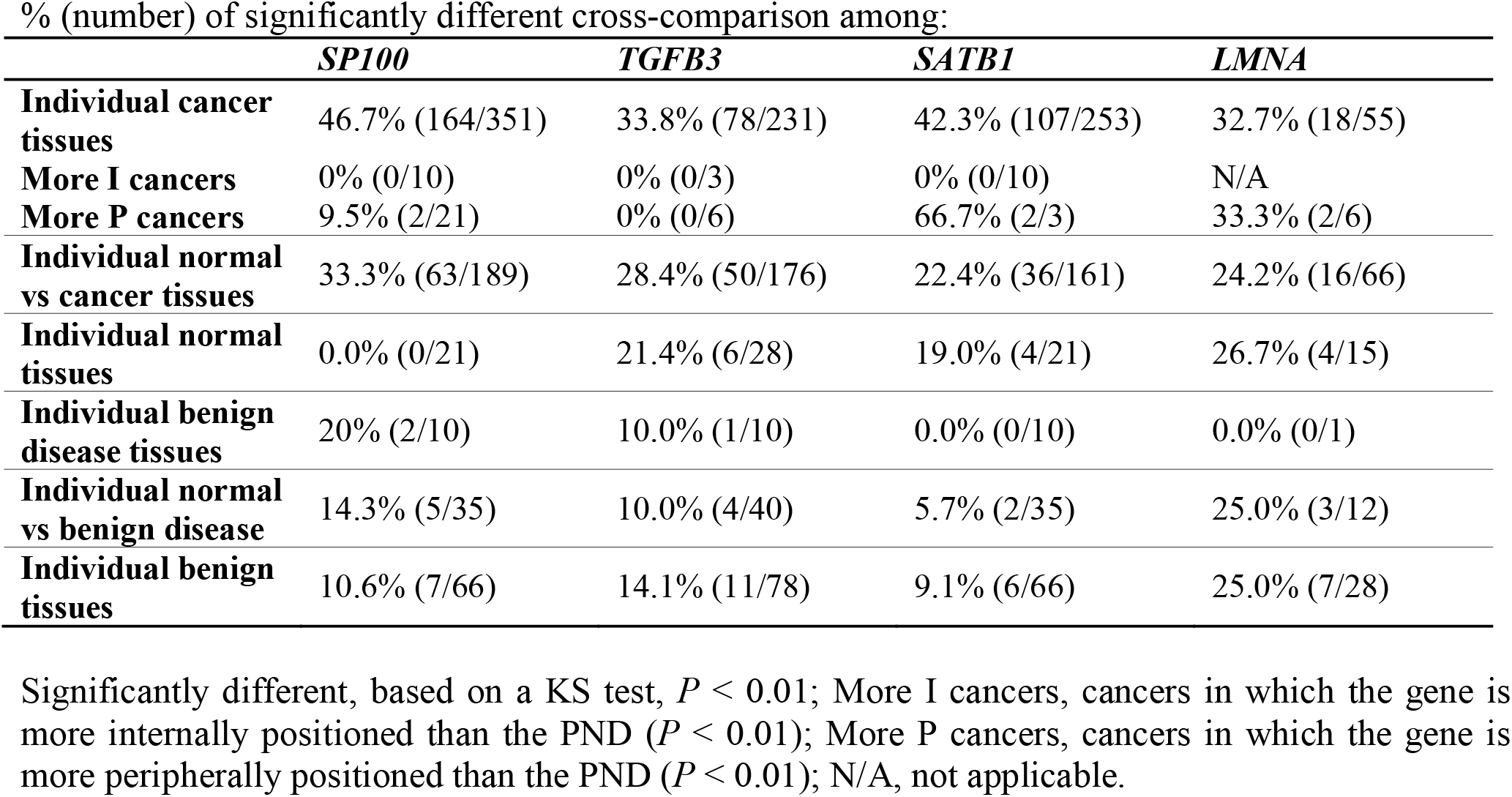
Cross-comparisons between individual tissues.

% (number) of significantly different cross-comparison among:
|  | *SP100* | *TGFB3* | *SATB1* | *LMNA* |
| --- | --- | --- | --- | --- |
| **Individual cancer tissues** | 46.7% (164/351) | 33.8% (78/231) | 42.3% (107/253) | 32.7% (18/55) |
| **More I cancers** | 0% (0/10) | 0% (0/3) | 0% (0/10) | N/A |
| **More P cancers** | 9.5% (2/21) | 0% (0/6) | 66.7% (2/3) | 33.3% (2/6) |
| **Individual normal vs cancer tissues** | 33.3% (63/189) | 28.4% (50/176) | 22.4% (36/161) | 24.2% (16/66) |
| **Individual normal tissues** | 0.0% (0/21) | 21.4% (6/28) | 19.0% (4/21) | 26.7% (4/15) |
| **Individual benign disease tissues** | 20% (2/10) | 10.0% (1/10) | 0.0% (0/10) | 0.0% (0/1) |
| **Individual normal vs benign disease** | 14.3% (5/35) | 10.0% (4/40) | 5.7% (2/35) | 25.0% (3/12) |
| **Individual benign tissues** | 10.6% (7/66) | 14.1% (11/78) | 9.1% (6/66) | 25.0% (7/28) |

Significantly different, based on a KS test, $P < 0.01$ ; More I cancers, cancers in which the gene is more internally positioned than the PND ( $P < 0.01$ ); More P cancers, cancers in which the gene is more peripherally positioned than the PND ( $P < 0.01$ ); N/A, not applicable.

### *SP100* and *TGFB3* Exhibit Differential Positioning Patterns Between Low and Intermediate/high Gleason Score Cancers

Next, we sought to determine if the differences in gene repositioning patterns between prostate cancers correlated with clinicopathological features. Comparing RRDs between individual cancer tissues was not useful for subgrouping cancers. For the most part, there was a similar proportion of cross-comparisons between cancers that were significantly different to each other within clinically relevant subgroups as there was between subgroups (for *P*-values see Supplementary Tables 2-5). For example, *SP100* was in a significantly different position in 49.1% (27/55) of cross-comparisons amongst Gleason score 2-6 prostate cancers, and 53.4% (47/88) of cross-comparisons when Gleason score 2-6 cancers were compared to Gleason score 7 cancers, and 50% (44/88) of cross-comparisons between Gleason score 2-6 and Gleason score 8-10 cancers (Supplementary Table 2).

In contrast, the behavior of a gene in a cancerous tissue compared to the PND was a better indicator to detect differential positioning between subgroups (Figure 1, Supplementary Figure 2, Tables 1, 3 and 4, Supplementary Tables 2-5). We first compared positioning patterns to Gleason score. In line with the clinical risk assessment of prostate cancers (Thompson et al., 2007), we classified Gleason scores of 2-6 as a low Gleason score, Gleason score 7 as intermediate, and scores of 8-10 as a high Gleason score. There was a modest increase in the proportion of cancer specimens with either *SATB1* or *LMNA* repositioned, compared to the PND, with increasing Gleason score, however, in the case of *LMNA* this may be due to the small sample size (Table 3). *SATB1* was in a statistically different nuclear position in 25% (2/8) of low Gleason score cancers, 33.3% (3/9) Gleason score 7 cancers and 50% (3/6) high Gleason score cancers. Similarly, *LMNA* repositioned in 33.3% of low (2/6) and intermediate (1/3) Gleason score cancers, and 50% (1/2) of high Gleason score cancers. Inclusion of the direction of repositioning did little to aid stratification (Table 3). Thus, we concluded that neither *SATB1* nor *LMNA* are biomarkers of Gleason score. The proportion of cancers in which *SP100* and *TGFB3* repositioned also did not stratify Gleason score groups. *SP100* was slightly more frequently repositioned in low Gleason score cancers specimens, repositioning in 54.5% (6/11) of low Gleason score cancers, 37.5% (3/8) of intermediate Gleason score cancers, and 37.5% (3/8) of high Gleason score cancers. *TGFB3* repositioned in 30% (3/10) low Gleason score cancer tissues, 20% (1/5) of intermediate Gleason score cancers, and 42.9% (3/7) of high Gleason score cancers. On the other hand, in cancer specimens in which either gene repositioned, the direction of repositioning correlated with Gleason score (Supplementary Figure 2A, Table 4). Both *SP100* and *TGFB3* shifted to a more peripheral position in 100% of the low Gleason score cancers in which these genes showed altered radial position. In contrast, *SP100* and *TGFB3* were in a more internal position in 83.3% (5/6) and 75.0% (3/4), respectively, of the Gleason score 7 and higher cancers in which they repositioned. Intermediate/high Gleason score cancer tissue repositioning is not exclusively more internal, since for both genes a more peripheral positioning was detected in a Gleason score 9 prostate cancer (Supplementary Figure 2A, Tables 1 and 4). The positioning patterns of *SP100* and *TGFB3* could not distinguish intermediate Gleason score cancer tissues from high Gleason score cancer tissues (Supplementary Figure 2A, Tables 1 and 4).

**Table 3.**
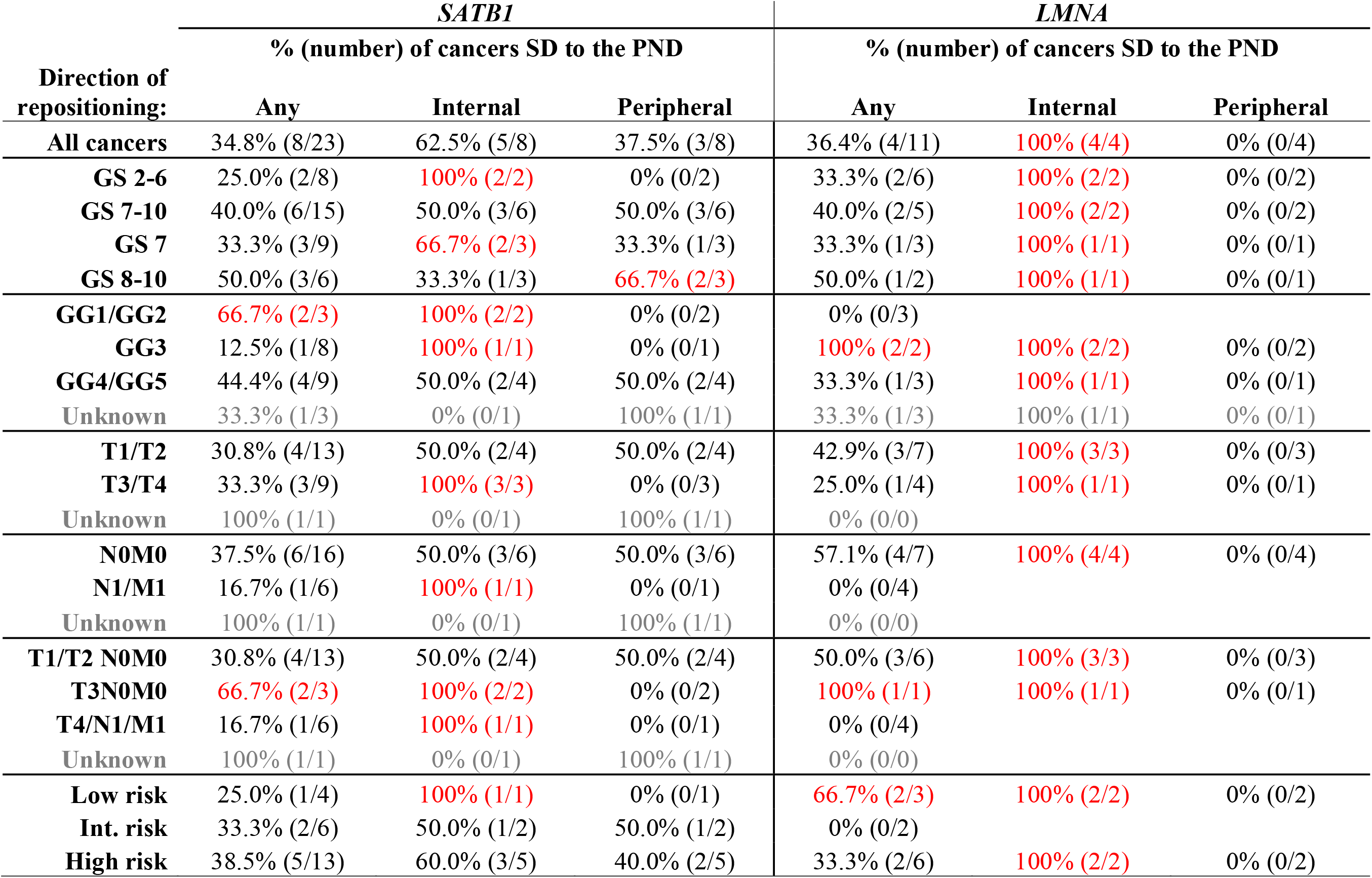
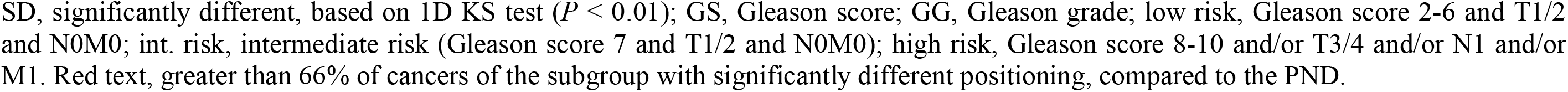
Positioning patterns for *SATB1* and *LMNA* by prostate cancer subgroups.

| Direction of repositioning: | *SATB1* |  |  | *LMNA* |  |  |
| --- | --- | --- | --- | --- | --- | --- |
|  | % (number) of cancers SD to the PND |  |  | % (number) of cancers SD to the PND |  |  |
|  | Any | Internal | Peripheral | Any | Internal | Peripheral |
| **All cancers** | 34.8% (8/23) | 62.5% (5/8) | 37.5% (3/8) | 36.4% (4/11) | 100% (4/4) | 0% (0/4) |
| **GS 2-6** | 25.0% (2/8) | 100% (2/2) | 0% (0/2) | 33.3% (2/6) | 100% (2/2) | 0% (0/2) |
| **GS 7-10** | 40.0% (6/15) | 50.0% (3/6) | 50.0% (3/6) | 40.0% (2/5) | 100% (2/2) | 0% (0/2) |
| **GS 7** | 33.3% (3/9) | 66.7% (2/3) | 33.3% (1/3) | 33.3% (1/3) | 100% (1/1) | 0% (0/1) |
| **GS 8-10** | 50.0% (3/6) | 33.3% (1/3) | 66.7% (2/3) | 50.0% (1/2) | 100% (1/1) | 0% (0/1) |
| **GG1/GG2** | 66.7% (2/3) | 100% (2/2) | 0% (0/2) | 0% (0/3) |  |  |
| **GG3** | 12.5% (1/8) | 100% (1/1) | 0% (0/1) | 100% (2/2) | 100% (2/2) | 0% (0/2) |
| **GG4/GG5** | 44.4% (4/9) | 50.0% (2/4) | 50.0% (2/4) | 33.3% (1/3) | 100% (1/1) | 0% (0/1) |
| **Unknown** | 33.3% (1/3) | 0% (0/1) | 100% (1/1) | 33.3% (1/3) | 100% (1/1) | 0% (0/1) |
| **T1/T2** | 30.8% (4/13) | 50.0% (2/4) | 50.0% (2/4) | 42.9% (3/7) | 100% (3/3) | 0% (0/3) |
| **T3/T4** | 33.3% (3/9) | 100% (3/3) | 0% (0/3) | 25.0% (1/4) | 100% (1/1) | 0% (0/1) |
| **Unknown** | 100% (1/1) | 0% (0/1) | 100% (1/1) | 0% (0/0) |  |  |
| **N0M0** | 37.5% (6/16) | 50.0% (3/6) | 50.0% (3/6) | 57.1% (4/7) | 100% (4/4) | 0% (0/4) |
| **N1/M1** | 16.7% (1/6) | 100% (1/1) | 0% (0/1) | 0% (0/4) |  |  |
| **Unknown** | 100% (1/1) | 0% (0/1) | 100% (1/1) | 0% (0/0) |  |  |
| **T1/T2 N0M0** | 30.8% (4/13) | 50.0% (2/4) | 50.0% (2/4) | 50.0% (3/6) | 100% (3/3) | 0% (0/3) |
| **T3N0M0** | 66.7% (2/3) | 100% (2/2) | 0% (0/2) | 100% (1/1) | 100% (1/1) | 0% (0/1) |
| **T4/N1/M1** | 16.7% (1/6) | 100% (1/1) | 0% (0/1) | 0% (0/4) |  |  |
| **Unknown** | 100% (1/1) | 0% (0/1) | 100% (1/1) | 0% (0/0) |  |  |
| **Low risk** | 25.0% (1/4) | 100% (1/1) | 0% (0/1) | 66.7% (2/3) | 100% (2/2) | 0% (0/2) |
| **Int. risk** | 33.3% (2/6) | 50.0% (1/2) | 50.0% (1/2) | 0% (0/2) |  |  |
| **High risk** | 38.5% (5/13) | 60.0% (3/5) | 40.0% (2/5) | 33.3% (2/6) | 100% (2/2) | 0% (0/2) |

SD, significantly different, based on 1D KS test ( $P < 0.01$ ); GS, Gleason score; GG, Gleason grade; low risk, Gleason score 2-6 and T1/2 and N0M0; int. risk, intermediate risk (Gleason score 7 and T1/2 and N0M0); high risk, Gleason score 8-10 and/or T3/4 and/or N1 and/or M1. Red text, greater than 66% of cancers of the subgroup with significantly different positioning, compared to the PND.

**Table 4.**
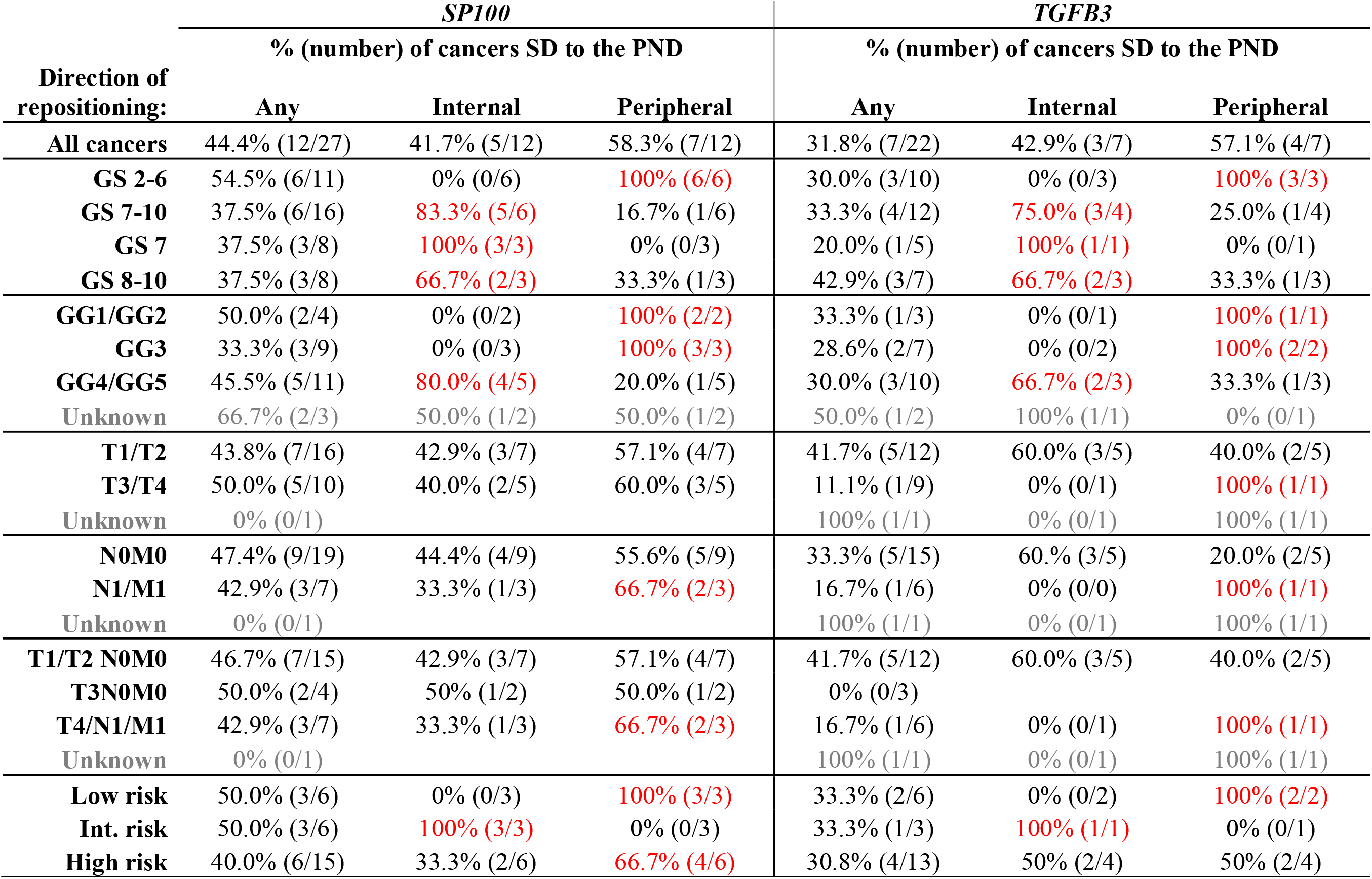
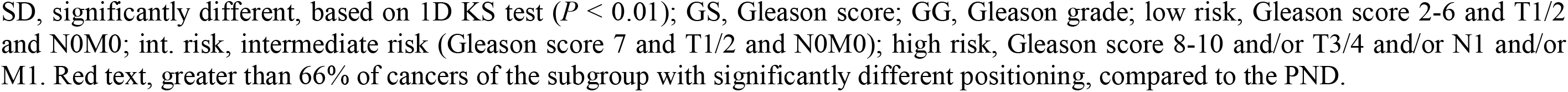
Positioning patterns for *SP100* and *TGFB3* by prostate cancer subgroups.

| Direction of repositioning: | *SP100* |  |  | *TGFB3* |  |  |
| --- | --- | --- | --- | --- | --- | --- |
|  | % (number) of cancers SD to the PND |  |  | % (number) of cancers SD to the PND |  |  |
|  | Any | Internal | Peripheral | Any | Internal | Peripheral |
| **All cancers** | 44.4% (12/27) | 41.7% (5/12) | 58.3% (7/12) | 31.8% (7/22) | 42.9% (3/7) | 57.1% (4/7) |
| **GS 2-6** | 54.5% (6/11) | 0% (0/6) | 100% (6/6) | 30.0% (3/10) | 0% (0/3) | 100% (3/3) |
| **GS 7-10** | 37.5% (6/16) | 83.3% (5/6) | 16.7% (1/6) | 33.3% (4/12) | 75.0% (3/4) | 25.0% (1/4) |
| **GS 7** | 37.5% (3/8) | 100% (3/3) | 0% (0/3) | 20.0% (1/5) | 100% (1/1) | 0% (0/1) |
| **GS 8-10** | 37.5% (3/8) | 66.7% (2/3) | 33.3% (1/3) | 42.9% (3/7) | 66.7% (2/3) | 33.3% (1/3) |
| **GG1/GG2** | 50.0% (2/4) | 0% (0/2) | 100% (2/2) | 33.3% (1/3) | 0% (0/1) | 100% (1/1) |
| **GG3** | 33.3% (3/9) | 0% (0/3) | 100% (3/3) | 28.6% (2/7) | 0% (0/2) | 100% (2/2) |
| **GG4/GG5** | 45.5% (5/11) | 80.0% (4/5) | 20.0% (1/5) | 30.0% (3/10) | 66.7% (2/3) | 33.3% (1/3) |
| **Unknown** | 66.7% (2/3) | 50.0% (1/2) | 50.0% (1/2) | 50.0% (1/2) | 100% (1/1) | 0% (0/1) |
| **T1/T2** | 43.8% (7/16) | 42.9% (3/7) | 57.1% (4/7) | 41.7% (5/12) | 60.0% (3/5) | 40.0% (2/5) |
| **T3/T4** | 50.0% (5/10) | 40.0% (2/5) | 60.0% (3/5) | 11.1% (1/9) | 0% (0/1) | 100% (1/1) |
| **Unknown** | 0% (0/1) |  |  | 100% (1/1) | 0% (0/1) | 100% (1/1) |
| **N0M0** | 47.4% (9/19) | 44.4% (4/9) | 55.6% (5/9) | 33.3% (5/15) | 60.0% (3/5) | 20.0% (2/5) |
| **N1/M1** | 42.9% (3/7) | 33.3% (1/3) | 66.7% (2/3) | 16.7% (1/6) | 0% (0/0) | 100% (1/1) |
| **Unknown** | 0% (0/1) |  |  | 100% (1/1) | 0% (0/1) | 100% (1/1) |
| **T1/T2 N0M0** | 46.7% (7/15) | 42.9% (3/7) | 57.1% (4/7) | 41.7% (5/12) | 60.0% (3/5) | 40.0% (2/5) |
| **T3N0M0** | 50.0% (2/4) | 50% (1/2) | 50.0% (1/2) | 0% (0/3) |  |  |
| **T4/N1/M1** | 42.9% (3/7) | 33.3% (1/3) | 66.7% (2/3) | 16.7% (1/6) | 0% (0/1) | 100% (1/1) |
| **Unknown** | 0% (0/1) |  |  | 100% (1/1) | 0% (0/1) | 100% (1/1) |
| **Low risk** | 50.0% (3/6) | 0% (0/3) | 100% (3/3) | 33.3% (2/6) | 0% (0/2) | 100% (2/2) |
| **Int. risk** | 50.0% (3/6) | 100% (3/3) | 0% (0/3) | 33.3% (1/3) | 100% (1/1) | 0% (0/1) |
| **High risk** | 40.0% (6/15) | 33.3% (2/6) | 66.7% (4/6) | 30.8% (4/13) | 50% (2/4) | 50% (2/4) |

SD, significantly different, based on 1D KS test ( $P < 0.01$ ); GS, Gleason score; GG, Gleason grade; low risk, Gleason score 2-6 and T1/2 and N0M0; int. risk, intermediate risk (Gleason score 7 and T1/2 and N0M0); high risk, Gleason score 8-10 and/or T3/4 and/or N1 and/or M1. Red text, greater than 66% of cancers of the subgroup with significantly different positioning, compared to the PND.

We also assessed if positioning patterns correlated with Gleason grade. For all four genes, increasing Gleason grade did not correlate with the percentage of cancer specimens in which the genes repositioned (Tables 3 and 4). Consistent with Gleason score, the direction that *SATB1* and *LMNA* repositioned did not aid in stratifying cancers by Gleason grade (Table 3), but the direction of repositioning did correlate with Gleason grade for *SP100* and *TGFB3* (Table 4). As with Gleason score, in the more highly differentiated cancers (Gleason grades 1-3) *SP100* and *TGFB3* repositioned towards the nuclear periphery in 100% (5 and 3 cancers, respectively) of the cancers in which these genes repositioned. Yet, in the poorly differentiated cancers (Gleason grade 4 and 5) both genes preferentially repositioned towards the nuclear interior. *SP100* was more internally positioned in 80% (4/5) of Gleason grade 4 and 5 cancer in which *SP100* repositioned and *TGFB3* was more internally positioned in 66.7% (2/3) of the Gleason grade 4 and 5 cancers in which *TGFB3* was repositioned, compared to the PND (Table 4). The similarity between Gleason grade and Gleason score positioning patterns are not surprising given that Gleason score is the sum of the two most prominent Gleason grades (Epstein et al., 2005;Epstein, 2018). An important caveat to be noted is that while the subgrouping of the cancers was based on the most predominant Gleason grade of the tissue, it is not necessarily the predominant Gleason grade of the nuclei analysis from each specimen.

Collectively, these observations demonstrate that while positioning patterns performed less well than the Gleason system at stratifying cancers, we identify differential gene positioning patterns between subgroups of prostate cancers.

### Multiplexing *SP100* and *TGFB3* Improves Detecting Intermediate and High Gleason Score Cancers

Although both *SP100* and *TGFB3* displayed differential positioning patterns between low and intermediate/high Gleason score cancer specimens, the sensitivity for subgrouping prostate cancers by Gleason score based on positioning patterns is low. Using a more peripheral positioning of *SP100* compared to the PND as a marker of low Gleason score cancers, the false negative rate (percentage of cancers without a more peripheral positioning) is 45.4% (5/11 low Gleason score cancers; Table 1). For *TGFB3* the false negative rate is even higher at 70% (7/10; Table 1). Additionally, using this criterion, false positive cancers were identified for both genes. More peripheral positioning was detected in one high Gleason score specimen for both *SP100* and *TGFB3*, resulting in a false positive rate of 6.3% (1/16) and 8.3% (1/12), respectively, for intermediate and high Gleason score cancers (Table 1). Neither gene was more internally positioned in low Gleason score cancers (Table 1). Using a more internal positioning pattern as a biomarker of intermediate and high Gleason score cancers resulted in a false negative rate of 62.5% (10/16) and 66.7% (8/12) for *SP100* and *TGFB3*, respectively (Table 1).

We have previously demonstrated that the sensitivity of diagnostic GPBs can be improved by multiplexing (Meaburn et al., 2009;Leshner et al., 2016). We therefore evaluated if combining positioning data from *SP100* and *TGFB3* would increase the number of cancers classified as low or intermediate/high Gleason score based on gene positioning patterns. Importantly for multiplexing to improve the sensitivity, *SP100* and *TGFB3* would need to be frequently repositioned in different cancer specimens. Of a subset of 19 cancer tissues in which both genes were positioned, 10 (52.6%) had differential repositioning patterns for *SP100* to that of *TGFB3* (Table 1, Supplementary Table 7). For six of these cancers *SP100* was repositioned but *TGFB3* was not, whereas in three cancers only *TGFB3* was repositioned. For one cancer sample, both genes were repositioned, compared to their PND, but they relocated in opposite directions, with *SP100* being more peripherally positioned, while *TGFB3* was more internally positioned (Table 1, Supplementary Table 7). However, multiplexing the two genes did not improve the sensitivity to detect low Gleason score cancers above using *SP100* alone. *SP100* was more peripherally positioned in all five cancers where at least one of the two genes repositioned (Table 1, Supplementary Table 7). Nevertheless, multiplexing increased the sensitivity of detecting intermediate/high Gleason score cancers (Table 1, Supplementary Table 7). At least one gene repositioned in eight of the 11 (72.7%) Gleason score 7 and higher cancers. Both *SP100* and *TGFB3* contributed to the increased proportion of cancer specimens with repositioning events. Of the seven cancers with only one of the two genes more internally repositioned, *SP100* was more internally repositioned in four and *TGFB3* more internally positioned in three cancers (Table 1, Supplementary Table 7). Using a more internal position of at least one of *SP100* or *TGFB3* the false negative rate for detecting intermediate or high Gleason score cancers was reduced to 36.4% (4/11). While most of the repositioning events in intermediate/high Gleason score cancers were to a more internal position, in one cancer the only repositioning event resulted in a more peripheral location of *TGFB3* and in another cancer tissue, there was both a more peripheral and more internal repositioning events, with *SP100* more peripherally position and *TGFB3* more internally located (Table 1, Supplementary Table 7).

Gleason score is not a perfect measure of risk. Given the variability in positioning patterns within the same Gleason group, we asked if the positioning patterns could be useful to distinguish aggressive low Gleason core cancers from non-aggressive low Gleason score cancers. Such a marker would aid treatment decisions. However, *SP100* and *TGFB3* were repositioned in a similar proportion of low Gleason score cancers with or without metastasis (Table 1). *SP100* repositioned in 50% (2/4) of low Gleason score cancers that had metastasized and 57.1% (4/7) of low Gleason score cancers without metastases. Likewise, *TGFB3* repositioned in 33.3% (1/3) of metastatic low Gleason score cancer specimens and 28.8% (2/7) of non-metastatic low Gleason score cancer specimens (Table 1). Thus, in addition to the high false negative rate for Gleason score, *SP100* and *TGFB3* can not distinguish aggressive low Gleason score cancers from non-aggressive low Gleason score cancers, limiting their clinical potential.

### Low Gleason Score Cancer Gene Positioning Patterns are Distinct from Benign Disease

Given the fact that low Gleason score cancers are fairly well differentiated tissues, it is possible that low Gleason score cancers have a similar genome organization to benign disease. We therefore sought to determine the cancer-specificity of the repositioning events. We positioned *SP100*, *TGFB3*, *SATB1* and *LMNA* in non-cancerous prostate tissues (Figure 2, Supplementary Figure 1, Tables 5 and 6, Supplementary Tables 2-5). For all four genes we found that the positioning patterns were highly similar between benign tissues. For *SP100, TGFB3* and *SATB1*, only 9.1%-14.1% of comparisons between the individual non-cancerous tissues reached significance. There was a little more variability between benign tissues for *LMNA*, where 25% of cross-comparisons between benign tissues were significantly different (Figure 2, Table 2, Supplementary Table 2-5). There was also little repositioning of the four genes in benign tissues when compared to the PND, with repositioning in 7.7%-16.7% of benign tissues, depending on the gene (Figure 2, Tables 5 and 6). *SP100* was statistically similarly positioned in all seven normal tissues, compared to the PND, but was significantly repositioned in 40% (2/5) of benign disease tissue (Figure 2, Tables 5 and 6). However, the positioning patterns of *SP100* were distinct in benign disease and low Gleason score cancer, since it was more internally localized in the two benign disease tissues, yet more peripherally located in low Gleason score cancers (Tables 1 and 5).

**Figure 2.**
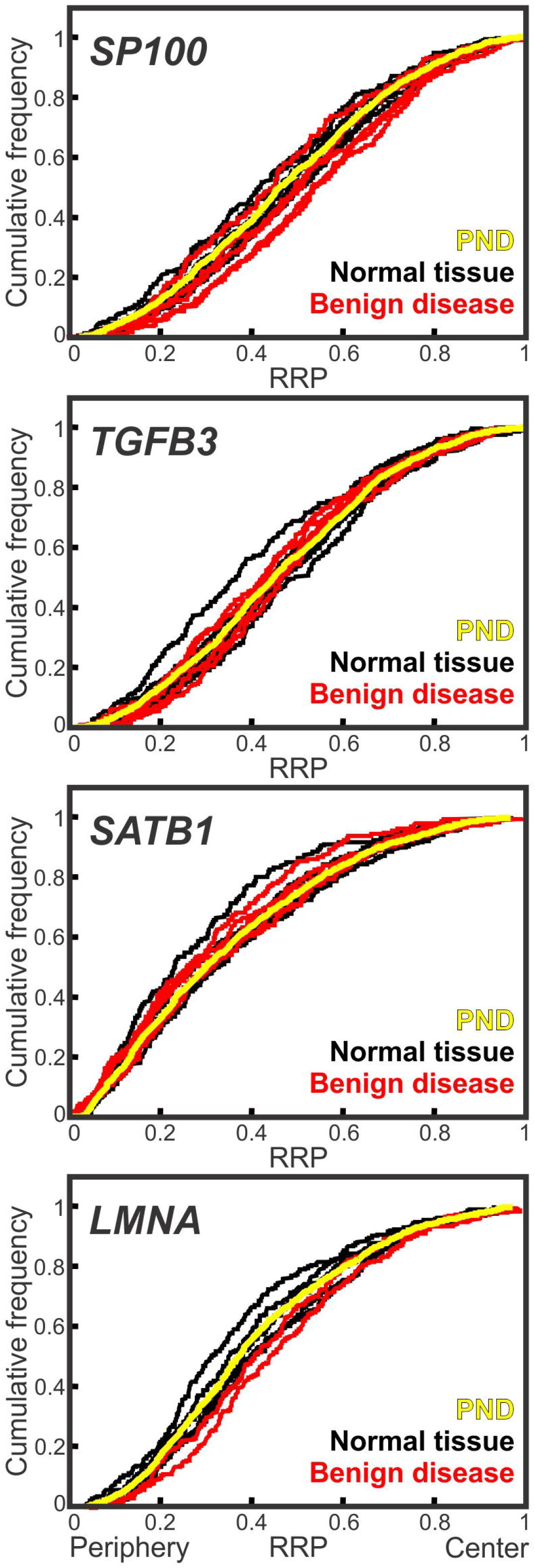
Conserved spatial organization of the genome in benign tissues. Cumulative RRDs for the indicated genes in normal prostate tissue (black), benign disease (red) and the pooled normal distribution (PND; yellow). RRP, relative radial position.

**Table 5.**
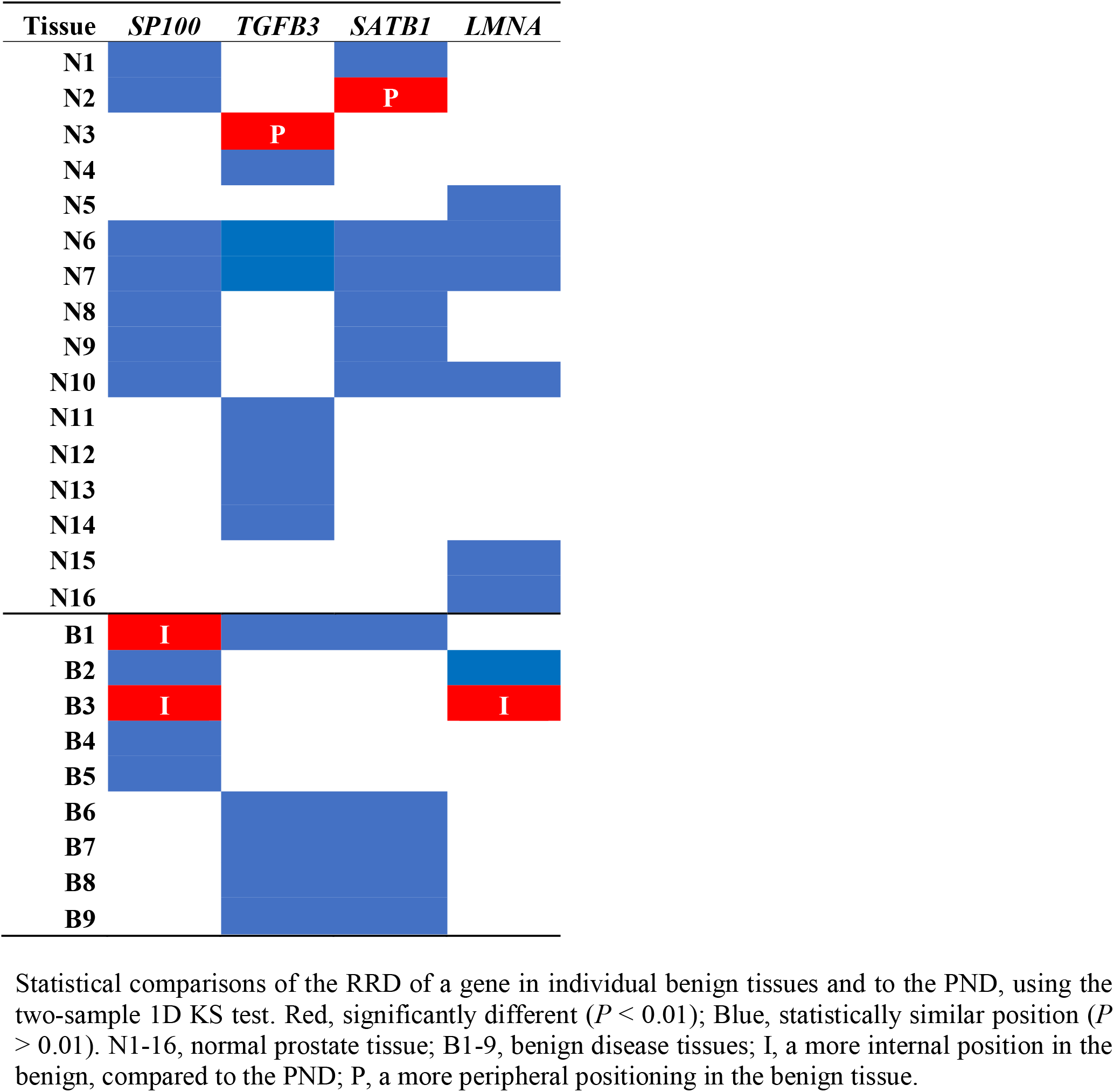
Conservation of positioning between normal prostate tissues and in benign disease.

**Table 6.** Comparison of individual benign tissue to the pooled normal.

|  | *SP100* | *TGFB3* | *SATB1* | *LMNA* |
| --- | --- | --- | --- | --- |
| Normal tissues | 0.0% (0/7) | 12.5% (1/8) | 14.3% (1/7) | 0.0% (0/6) |
| Benign disease | 40.0 (2/5) | 0.0% (0/5) | 0.0% (0/5) | 50.0% (1/2) |
| Total benign tissues | 16.7% (2/12) | 7.7% (1/13) | 8.3% (1/12) | 12.5% (1/8) |

% (number) of benign tissues where the RRD is significantly different to the PND ( $P < 0.01$ ; KS test).

Inclusion of the direction of repositioning in the analysis further confirmed the specificity of the repositioning events to the different Gleason score subgroups. When using more peripheral positioning, compared to the PND, as a marker of low Gleason score prostate cancer, the false positive rate for *SP100* is very low, at 3.6%, since it is more peripherally positioned in only one of 28 normal, benign disease and higher Gleason score cancer tissues (Tables 1 and 5). Similarly, *TGFB3* was repositioned in a single normal tissue (12.5%; 1/8) and in none of the benign disease tissues (0/5), compared to the PND (Figure 2, Tables 5 and 6). Unlike *SP100*, the direction *TGFB3* reposition in the normal tissue was the same as in low Gleason score cancer. However, the false positive rate for using a more peripheral positioning of *TGFB3* was relatively low at 8%, because it was more peripherally positioned in two of the 25 benign tissues and higher Gleason score cancers (Tables 1 and 5). The false positive rate of using a more internal position of *SP100* or *TGFB3* to detect intermediate/ high Gleason score cancers is also low, at 8.7% and 0%, respectively, since *SP100* was more internally repositioned in only two of the 23 benign tissues and low Gleason score cancer specimens and *TGFB3* was not more internally repositioned in these groups of tissues (*N*=23; Table 1 and 5). Taken together, we find the spatial organization of the genome is generally conserved between benign tissues, and benign disease tissue have a distinct genome organization to both low and intermediate/high Gleason score cancers.

### *LMNA* Repositions in Low Risk and Non-Metastatic Cancers

Having determined the biomarker potential of the candidate genes for subgrouping prostate cancers by Gleason score, we compared their positioning patterns to other clinical markers of poor patient outcome. The TNM staging system is commonly used to aid the prediction of the aggressiveness of the cancer and the risk of poor patient outcome (Thompson et al., 2007). In stage T1 and T2 prostate cancers the tumor is contained within the prostate, whereas in stages T3 and T4 the cancer has spread from the prostate into the surrounding tissue. T1 and T2 cancers are lower risk cancers and respond better to treatment than T3 and T4 prostate cancers (Thompson et al., 2007;Chang et al., 2014). The N and M score indicates whether the cancer has spread beyond the surrounding tissue. In N0 cancers, no cancer cells are detected in the regional lymph nodes, whereas N1 denotes that the cancer has spread into the regional lymph nodes. For M0 cancers, no distant metastasis are detected, while distant metastases, to non-regional lymph nodes or organs, have occurred in M1 cancers (Thompson et al., 2007).

Positioning patterns for *SP100* and *SATB1* were similar in low and high T stage cancer specimens and could not be used to distinguish the different T stage group cancers from each other (Tables 3 and 4). *TGFB3* and *LMNA* were both more frequently repositioned in low T stage cancers to that of high T stage cancers, but with high false positive rates (Tables 3 and 4). *TGFB3* repositioned in 41.7% (5/12) of T1/2 cancers, 11.1% (1/9) of T3/4 cancers, and 7.7% (1/13) of benign tissues, compared to the PND (Tables 1, 4 and 6). This equates to a false negative rate of 58.3% (7/12) and a false positive rate of 9.1% (2/22) for using the reposition of *TGFB3* to detect low T stage prostate cancer. Similarly, *LMNA* repositioned in 42.9% (3/7) of low T stage cancer, 25% (1/4) of high T stage cancers and 12.5% (1/8) of benign tissues (Tables 3 and 6), making the false negative and positive rates of using the repositioning of *LMNA* to demark low T stage cancers 57.1% (4/7) and 16.7% (2/12), respectively.

Clinically, multiple factors are combined to determine risk of poor outcome. Therefore, we also compared gene positioning patterns with a multifactorial determinant of risk using standard clinical risk assessment criteria (Thompson et al., 2007;Chang et al., 2014) with the exception of PSA levels, since no information on serum PSA were available for our specimens. Moreover, we included N1 and/or M1 cancers in the high-risk group, since they are known aggressive cancers. We classified low risk cancer as Gleason score 2-6 and T1/2 and N0M0 cancers; intermediate risk cancers as Gleason score 7 and T1/2 and N0M0; high risk as Gleason score 8-10 and/or T3/4 and/or N1 and/or M1 prostate cancers. The positioning patterns of *SP100*, *TGFB3* and *SATB1* were similar in all three risk groups, and thus could not be used as markers of risk (Tables 3 and 4). Conversely, *LMNA* was more frequently repositioned in low risk prostate cancer, since it repositioned in 66.7% (2/3) of low risk cancers, 0% (0/2) of intermediate risk and 33.3% (2/6) of high-risk groups (Table 3).

Finally, as a more direct measure of the aggressiveness of a cancer we compared non-metastatic cancers (N0M0) and metastatic (N1/M1) prostate cancers (Supplementary Figure 2B, Table 1, 3 and 4). *SP100* repositioned, compared to the PND, in a similar proportion of N0M0 (47.4%; 9/19) and metastatic (42.9%; 3/7) cancer specimens. Furthermore, the direction of repositioning was similarly mixed in both groups of cancer (Table 4). The remaining three genes repositioned more frequently in non-metastatic cancers. For *TGFB3* and *SATB1* this difference is small, with the genes repositioned in ~33.3% (5/15 and 6/16 respectively; false negative rate ~66.7%) of non-metastatic cancers and 16.7% (1/6) of metastatic cancers (Table 3 and 4). *LMNA* was the best marker of non-metastatic cancers. *LMNA* repositioned in 57.1% (4/7) of non-metastatic prostate cancer specimens and was not repositioned in metastatic (0/4) cancer tissues (Supplementary Figure 2B, Table 3). As with *SP100* and *TGFB3* as markers of Gleason score, the false negative rate for using *LMNA* positioning as a marker of non-metastatic cancer was high at 42.9% (3/7), and the false positive rate is relatively low at 8.3% (1/12) (Tables 1, 3 and 5).

Taken together, our data suggest that there are distinct spatial gene positioning patterns between some subgroups of prostate cancers, although the false negative rates were generally high, limiting their potential for clinical use.

## Discussion

To reduce overtreatment in cancer patients that receive no benefit from medical intervention, there is an urgent need for biomarkers that predict the aggressiveness of a cancer. Here, we assess the feasibility of utilizing the spatial positioning patterns of genes in interphase nuclei for prognostic purposes in prostate cancer. We find a differential enrichment of specific positioning patterns for multiple genes between clinically relevant subgroups of prostate cancers. While the false positive rates for prognostic evaluation are low, the false negative rates are generally high, limiting clinical usefulness. Our results of subtype-specific genome organization patterns suggest that it should be possible to find clinically valuable prognostic GPB by screening additional genes and combinations of genes.

The spatial organization of the genome is altered in diseased cells, and at least some of the changes to genomic spatial positioning patterns are disease-specific (Meaburn, 2016). For instance, *HES5* repositions in breast cancer, but not in benign breast disease nor prostate cancer (Meaburn et al., 2009;Meaburn et al., 2016). Alternative spatial positioning patterns are not only found in cancers formed in different organs, but there is also heterogeneity in the spatial organization of the genome between individual cancers of the same type (Meaburn et al., 2009;Knecht et al., 2012;Leshner et al., 2016;Meaburn et al., 2016). We hypothesized that heterogeneity within a cancer type may reflect the aggressiveness of a cancer and therefore be of prognostic value. Prognosis-related repositioning of genomic regions in several types of cancer has previously been reported, with increased clustering of telomeres linked to poorer patient outcomes (Mai, 2018). For example, at the time of diagnosis telomeres were more likely to cluster in Hodgkin lymphoma patients whose disease later relapsed or progressed compared to patients who responded well to treatment (Knecht et al., 2012). Currently, Gleason score and the presence or absence of metastasis are key clinicopathological tumor features for predicting the aggressiveness of a prostate cancer. We find that *SP100* and *TGFB3* occupy alterative positions in low Gleason score cancers compared to higher Gleason scored cancers, and that *LMNA* repositions more internally in many non-metastatic and low risk prostate cancers, but infrequently reposition in high risk/aggressive cancers.

Our previous identification of diagnostic GPBs was based on the percentage of cancer specimens in which a gene had an alternative radial position, compared to its distribution in normal tissues (Meaburn, 2016). Interestingly, for *SP100* and *TGFB3* it was not the repositioning itself, but the direction of repositioning that was useful for stratification of prostate cancers. Repositioning of either *SP100* or *TGFB3* towards the nuclear periphery was associated with low Gleason score whereas repositioning towards the interior was a marker of higher Gleason score cancers. The repositioning patterns of *SP100* and *TGFB3* could not distinguish intermediate from high Gleason score cancers. Nevertheless, this does not necessarily rule them out as useful clinical markers since low Gleason score cancers are less likely to receive treatment than intermediate or high Gleason score cancers (Thompson et al., 2007;Cooperberg et al., 2010;Leshner et al., 2016). Given that there is inter-and intra-observer variability when scoring cancers (Montironi et al., 2005), additional markers that can clarify if a cancer is Gleason score 6 (low) or 7 (intermediate) would be useful in guiding therapeutic choices. However, because the positioning patterns of our genes could not separate aggressive, metastatic low Gleason score cancers from non-metastatic low Gleason score cancers, they are unlikely to aid the decision of whether to treat a cancer or not. In keeping with a differential spatial genome organization in cancers above and below the treatment threshold, we previously found *MMP9* to reposition in 20% of low Gleason score cancers compared to 82% of intermediate/high Gleason score cancers (Leshner et al., 2016). Unlike *SP100* and *TGFB3*, the direction *MMP9* repositioned did not aid stratification (Leshner et al., 2016), unpublished data). *MMP9* was positioned predominantly in Gleason score 6 and 7 prostate cancers, making it unclear how specific these positioning patterns are more generally to the different Gleason scores subgroups. *SP100*, *TGFB3* and *MMP9* each map to different chromosomes (HSA 2, 14 and 20, respectively) and therefore represent independent repositioning events within the subgroups of different Gleason score cancers.

In our analysis we find low false positive rates for distinguishing low from intermediate/high Gleason score cancers. In keeping with previous studies (Borden and Manuelidis, 1988;Meaburn et al., 2009;Leshner et al., 2016;Meaburn et al., 2016), we find similar positioning patterns for both *SP100* and *TGFB3* amongst normal tissues and between normal and benign disease tissues, highlighting that the gene repositioning in cancer tissues were specific to cancer. Despite the fact that low Gleason score cancers represents fairly well differentiated tissues there were distinct positioning patterns for *SP100* and *TGFB3* between low Gleason score cancers and benign disease, which are considered differentiated tissues. *TGFB3* did not reposition in benign disease and *SP100* was repositioned in only 20% of the benign disease tissues. However, unlike low Gleason score cancers, *SP100* was more internally positioned in benign disease tissues, and therefore does not contribute to the false positive rate when using more peripheral positioning of these genes to detect low Gleason score cancers. Unlike biomarkers used to diagnose cancer, the false positive rate of detecting a subtype of cancer for prognostic purposes is not only generated from non-cancerous tissues, it needs to also include cancers from the alternative subgroups. Even so, the false positive rates of detecting low Gleason score prostate cancers, were low because the direction of repositioning for *SP100* and *TGFB3* was mostly specific to the subgroups. In contrast to the false positive rates, the false negative rates for *SP100* and *TGFB3* were high, at 45-70%. We have previously found that for some genes multiplexing reduces the false negative rate and thus the sensitivity of detecting cancer (Meaburn et al., 2009;Leshner et al., 2016). Constant with this, we find that multiplexing *SP100* and *TGFB3* reduces the false negative rate of detecting intermediate and higher Gleason score cancers. However, multiplexing with a more peripheral position of either *SP100* or *TGFB3* did not reduce the false negative rate for low Gleason score cancers from that of using *SP100* alone. We conclude that the observed high false negative rates reduce marker strength and the utility of these genes for prognostic purposes.

Even though the positioning patterns of *SP100* and *TGFB3* are inferior to the Gleason system at stratifying cancers, our results reveal subtype-specific genome organization. Similarly, the repositioning of *LMNA* is also subtype-specific, but in this case the repositioning occurs only in non-metastatic cancers, although also with a high false negative rate. Interestingly, the reorganization events between the different subtypes of prostate cancer appear to be gene-specific. *LMNA* and *SATB1* positioning patterns were not able to stratify prostate cancers by Gleason score and *SP100*, *TGFB3* and *SATB1* were not accurate markers of risk or aggressiveness of the cancer. Consistently, the radial repositioning patterns of *FLI1*, *MMP9* and *MMP2* also do not correlate with the risk/aggressiveness of prostate cancer (Leshner et al., 2016). Given that it can take many years after the initial diagnosis of prostate cancer to progress to recurrence, metastasis and/or lethality (Albertsen et al., 1998;D’Amico et al., 1998;Pound et al., 1999;Cooperberg et al., 2009), it will be necessary to analyze specimens with long-term (15+ years) follow-up to accurately assess the potential of spatial positioning for assessment of risk or aggressiveness.

It is unknown what mechanisms lead to the reorganization of the genome in disease, and many processes have been implicated in regulating spatial positioning patterns, including changes in gene expression, replication timing, chromatin modifications, altered amounts of nuclear proteins, making it likely that the mis-regulations of these cellular functions in cancer cells is related to the spatial mis-organization of the genome (Zink et al., 2004;Meaburn, 2016;Flavahan et al., 2017). The four genes we studied have all been associated with carcinogenesis and have a range of functions. SP100 is a major component of the PML nuclear body and has been implicated in transcription regulation, cellular stress, oxidative stress, telomere length and stability, senescence, apoptosis and DNA damage repair (Lallemand-Breitenbach and de The, 2010). However, most of the evidence for PML bodies role in cancer relates to the PML protein, not SP100, which has not been implicated in prostate cancer. TGFB3 is a cytokine, with important roles in development, wound healing, the immune response and acts as a tumor suppresser in early cancers but can switch to promoting cancer progression in later stages (Massague, 2008;Laverty et al., 2009). *TGFB3* gene expression levels have been identified as a potential biomarker for prostate cancer, being expressed at lower levels in prostate cancer than normal tissue (Wang et al., 2017). Moreover, *TGFB3* expression levels correlated weakly with both progression-free survival and Gleason score (Wang et al., 2017). SATB1, a nuclear architectural protein that facilitates DNA loop formation and chromatin remodeling (Kohwi-Shigematsu et al., 2013), promotes the progression of many cancers, including prostate cancer, and is overexpressed in high Gleason score cancers compared to low Gleason score cancers and in metastatic compared to non-metastatic prostate cancers (Mao et al., 2013;Shukla et al., 2013;Naik and Galande, 2019). *LMNA* encodes for A-type lamins, proteins that reside predominantly at the nuclear envelope, and have a variety of roles including in nuclear structure, transcription regulation, and spatial genome organization (Dittmer and Misteli, 2011). A-type lamins levels are altered in many types of cancer, with reduced levels often, but not always, linked to a tendency for a poorer prognostic outcome (Meaburn, 2016). It is not currently clear what effect prostate cancer has on A-type lamin protein level. On the one hand, levels of A-type lamns in prostate cancer have been correlated with poor outcome, with reduced levels associated with an increased risk of lymph node metastasis, and poor outcome in Gleason score 7 and higher prostate cancers (Saarinen et al., 2015). On the other hand, reduced A-type lamin levels in Gleason score 6 cancer compared to high Gleason score cancer, increased A-type lamin levels in cells at the invasive leading edge of prostate cancers, and enhanced migration and invasion in the presence of high A-type lamin levels have been reported (Skvortsov et al., 2011;Kong et al., 2012).

Increased cell proliferation is associated with a poor outcome for prostate cancer patients (Berlin et al., 2017), and several genomic loci are differentially positioned between proliferating and non-proliferating cells (Bridger et al., 2000;Meaburn and Misteli, 2008). However, variations in proliferation rate is unlikely to be a major determinant in the gene repositioning we detect. In fact, the vast majority of cells in a prostate cancer tumor are non-proliferating, with a mean of just 6.1% proliferating cells per cancer (Berlin et al., 2017). Furthermore, in a cell culture model of breast cancer, there were distinct genome spatial rearrangements associated with proliferation status to that of carcinogenesis (Meaburn and Misteli, 2008). Similarly, although we find that changes in copy number did not correlated with propensity to reposition (Supplementary Table 6; (Meaburn et al., 2009;Leshner et al., 2016;Meaburn et al., 2016), we can not fully rule out that structural genomic alterations have not influenced the spatial position of any of the genes in the tissues analyzed since, in some cases, genomic instability can lead to spatial reorganization of the genome (Croft et al., 1999;Taslerova et al., 2003;Harewood et al., 2010;Federico et al., 2019). Regardless, importantly for a clinical test, we find that even in the background of genomic instability it is still possible to use gene positioning to distinguish normal tissue from cancer (Meaburn et al., 2009;Leshner et al., 2016;Meaburn et al., 2016) and to stratify cancers into clinically distinct subgroups, as demonstrated in this study.

Taken together, this study assesses the utility of spatial gene positioning in the stratification of prostate cancers. Our results reveal correlations between gene location and the aggressiveness of a tumor, which may serve as the foundation for prognostic usage of gene positioning. While the genes analyzed here have a relatively high false negative rate of detecting cancer subgroups, our results encourage the exploration of additional candidate genes in larger sample sets for the discovery of spatial genome positioning patterns as prognostic biomarkers.

## Supporting information

Sub Table 3

Sub Table 5

## Author Contributions Statement

KM and TM conceived the study and wrote the manuscript. KM designed the study, performed the experiments, analyzed and interpreted data, and made the figures.

## Abbreviations

BAC: bacterial artificial chromosome
EDT: Euclidean distance transform
FISH: fluorescence *in situ* hybridization
FFPE: formalin-fixed paraffin embedded
GPB: gene positioning biomarker
HSA: human chromosome
KS test: Kolmogorov-Smirnov test
NAT: normal adjacent to tumor
PND: pooled normal distribution
PSA: serum prostate specific antigen
RRD: relative radial distribution
TMA: tissue microarray

## Acknowledgments

We thank Lawrence True for invaluable and insightful prostate pathology discussions, Prabhakar Gudla and Stephen Lockett for the FISH positioning analysis software; Delft University (Netherlands) for providing the DIPImage and PRTools toolboxes and Tatiana Karpova for microscopy support. Fluorescence imaging was performed at the National Cancer Institute (NCI) Fluorescence Imaging Facility.

