## Supplementary material for "Assessment of the Utility of Gene Positioning Biomarkers in the Stratification of Prostate Cancers": Sup Table 1

**Supplementary Table 1 Characterization of human prostate tissues**

**A:** Prostate cancer specimens

| **Sample ID** | **Gleason Score** | **Gleason Grade** | **TNM** | **Risk/aggressive** | **Grade** | **Stage** | **Age** | **Source** | **TMA** | **TMA Core/source ID** |
| --- | --- | --- | --- | --- | --- | --- | --- | --- | --- | --- |
| **C1** | 9 (4+5) | 4 | T3N0M1 | High | 3 | IV | 64 | Biomax | PR483c | C7 |
| **C2** | 9 (5+4) | 4-5 | T2N1M1c | High |  | IV | 75 | Biomax | PR243b | C2 |
| **C3** | 9 (5+4) | 5 | T3aN0M0 | High | 3 | III | 66 | Biomax | T195e | A4 |
| **C4** | 8 (4+4) | 4 | T3N0M0 | High | 2-3 | III | 70 | Biomax | T195e | B6 |
| **C5** | 9 (4+5) |  | T2N0M0 | High | III |  | 47 | Biomax | PRC481 | E3 |
| **C6** | 9 (4+5) |  | T2N0M0 | High | III |  | 77 | Biomax | PRC481 | F5 |
| **C7** | 9 (4+5) |  | T2N0M0 | High | III |  | 72 | Biomax | PRC481 | E2/F2 |
| **C8** | 8 (4+4) | 4 | Unknown | High | 2 |  | 82 | Biomax | PR808a | B2 |
| **C9** | 7 (3+4) | 4 | T3N0M1b | High | 2 | IV | 64 | Biomax | PR242b | C2 |
| **C10** | 7 (3+4) | 3 | T4N1M1c | High | 2 | IV | 74 | Biomax | PR483c | A1 |
| **C11** | 7 | 3 | T3N0 | High |  |  | 60's | L True | N/A | UW10 |
| **C12** | 7 | 3 | T2N0 | Intermediate |  |  | 50's | L True | N/A | UW9 |
| **C13** | 7 | 3 | T2N0 | Intermediate |  |  | 60's | L True | N/A | UW11 |
| **C14** | 7 (3+4) |  | T2N0M0 | Intermediate | III |  | 72 | Biomax | PRC481 | F7/E7 |
| **C15** | 7 (3+4) | 4 | T2N0M0 | Intermediate | 2 | IIA | 57 | Biomax | PR808a | B9 |
| **C16** | 7 (3+4) | 4 | T2N0M0 | Intermediate | 2 | IIA | 73 | Biomax | PR808a | D8 |
| **C17** | 7 (3+4) |  | T1N0M0 | Intermediate | II |  | 60 | Biomax | PRC481 | D4/C4 |
| **C18** | 7 |  | T2N0 | Intermediate |  |  | 50's | L True | N/A | UW12 |
| **C19** | 7 |  | T2N0 | Intermediate |  |  | 50's | L True | N/A | UW13 |
| **C20** | 4 (2+2) | 2 | T4N1M1 | High | 1 | IV | 75 | Biomax | PR242a/b | B2 |
| **C21** | 3 (1+2) | 2 | T3N1M1 | High | 1 | IV | 66 | Biomax | PR242a/b | A1/2 |
| **C22** | 6 (2+4) | 4 | T3N0M1b | High |  | IV | 64 | Biomax | PR242a | C2 |
| **C23** | 6 (3+3) | 3 | T3N1M0 | High | 2 | IV | 60 | Biomax | PR808a | A7/A8 |
| **C24** | 6 (3+3) | 3 | T3N0M0 | High | 2 | III | 62 | Biomax | PR483c | B8 |
| **C25** | 6 (3+3) | 3 | T2N0M0 | Low |  | II | 73 | Biomax | N/A | HuCAT371 |
| **C26** | 6 (3+3) | 3 | T2N0M0 | Low |  | II | 58 | Biomax | PR242a | B5 |
| **C27** | 6 |  | T2N0 | Low |  |  | 50's | L True | N/A | UW5 |
| **C28** | 5 (1+4) |  | T2N0M0 | Low | I-II |  | 61 | Biomax | PRC481 | D2/C2 |
| **C29** | 5 (2+3) | 3 | T2N0M0 | Low | 1 | I | 68 | Biomax | PR808a | A1/A2 |
| **C30** | 5 (2+3) | 3 | T2N0M0 | Low | 1 | I | 75 | Biomax | PR808a | B8 |
| **C31** | 4 (2+2) | 2 | T2N0M0 | Low | 1 | II | 71 | Biomax | PR483c | A5 |
| **C32** | 3 (1+2) | 1 | T1N0M0 | Low |  | I | 57 | Biomax | T195c | A5 |

**B:** Benign prostate tissues

| **Sample ID** | **Pathology** | **Age** | **Source** | **TMA** | **Core/source ID** |
| --- | --- | --- | --- | --- | --- |
| **N1** | Normal, absence ca | 35 | Biomax | N/A | HuFPT143 |
| **N2** | Normal, absence ca |  | L True | N/A | UW21 |
| **N3** | Normal, absence ca | 28 | Biomax | N/A | HuFPT141 |
| **N4** | Normal, absence ca | 54 | Imegenex | N/A | IMH-1035 |
| **N5** | Normal, absence ca |  | L True | N/A | UW23 |
| **N6** | NAT | 27 | Biomax | PR242a | D4 |
| **N7** | NAT | 25 | Biomax | PR242a | D5 |
| **N8** | NAT | 60's | L True | N/A | UW11 |
| **N9** | NAT | 50's | L True | N/A | UW9 |
| **N10** | NAT | 60's | L True | N/A | UW10 |
| **N11** | NAT |  | L True | N/A | UW7 |
| **N12** | NAT |  | L True | N/A | UW8 |
| **N13** | NAT | 50's | L True | N/A | UW18 |
| **N14** | NAT | 64 | Biochain | N/A | T2234201 |
| **N15** | NAT | 50's | L True | N/A | UW12 |
| **N16** | NAT | 50's | L True | N/A | UW13 |
| **B1** | Chronic prostatitis (inflammation) | 54 | Biomax | PRC481 | B3 |
| **B2** | Hyperplasia | 27 | Biomax | PR243b | B3 |
| **B3** | Hyperplasia | 35 | Biomax | PR243b | B7 |
| **B4** | Hyperplasia | 84 | Biomax | PRC481 | A8 |
| **B5** | Hyperplasia | 64 | Biomax | PRC481 | B7 |
| **B6** | Hyperplasia | 65 | Biomax | PRC481 | B5/A5 |
| **B7** | Hyperplasia | 65 | Biomax | PRC481 | B6 |
| **B8** | Hyperplasia | 68 | Biomax | PRC481 | F1/E1 |
| **B9** | Hyperplasia |  | L True | N/A | UW20 |

Source ID, deidentified specimen identification code from the source of the tissue. Biomax, US Biomax Inc; L.D.T., Dr. Lawrence True; Imgenex, Imgenex corporation; NAT, Normal Adjacent to Tumor; Normal, Normal tissue taken from cancer free prostates; TMA, Tissue microarray. Red: mark of aggressive/high risk cancer. Blue: mark of low risk cancer. Purple, intermediate Gleason score. M1b: metastasis in bone, M1c: Metastasis in another part of the body, with or without bone metastasis. N/A not applicable, single tissue slide. Grey sample ID: RRD’s for these tissues previously published (*SP100*, *SATB1*, *LMNA*: Leshner et al., 2016; *TGFB3*: Meaburn et al., 2016), compared to an updated PND (see Materials and Methods).
