## Supplementary material for "Assessment of the Utility of Gene Positioning Biomarkers in the Stratification of Prostate Cancers": Sup Table 7

**Supplementary Table 7: Multiplexing *SP100* and *TGFB3***

| **Sample ID** | ***SP100*** | ***TGFB3*** | **Gleason Score** | **TNM** |
| --- | --- | --- | --- | --- |
| **C1** | **I** |  | **9 (4+5)** | **T3N0M1** |
| **C3** |  |  | **9 (5+4)** | **T3AN0M0** |
| **C4** | **I** |  | **8 (4+4)** | **T3N0M0** |
| **C5** |  |  | **9 (4+5)** | **T2N0M0** |
| **C6** | **P** | **I** | **9 (4+5)** | **T2N0M0** |
| **C7** |  | **I** | **9 (4+5)** | **T2N0M0** |
| **C8** |  | **P** | **8 (4+4)** | **unknown** |
| **C10** |  |  | **7 (3+4)** | **T4N1M1c** |
| **C14** |  | **I** | **7 (3+4)** | **T2N0M0** |
| **C15** | **I** |  | **7 (3+4)** | **T2N0M0** |
| **C16** | **I** |  | **7 (3+4)** | **T2N0M0** |
| **C20** | **P** |  | **4 (2+2)** | **T4N1M1** |
| **C21** | **P** | **P** | **3 (1+2)** | **T3N1M1** |
| **C23** |  |  | **6 (3+3)** | **T3N1M0** |
| **C24** | **P** |  | **6 (3+3)** | **T3N0M0** |
| **C25** | **P** | **P** | **6 (3+3)** | **T2N0M0** |
| **C26** | **P** | **P** | **6 (3+3)** | **T2N0M0** |
| **C30** |  |  | **5 (2+3)** | **T2N0M0** |
| **C31** |  |  | **4 (2+2)** | **T2N0M0** |

Positioning patterns of *SP100* and *TGFB3* in cancer tissues (C1-C31), compared to the PND. Only cancers with both genes positioned shown (see Table 1). I denotes a more internal position in the cancer compared to the PND, P denotes a more peripheral position. Red boxes indicate gene is repositioned compared to the PND (*P* < 0.01; KS test). Blue boxes indicate similar radial position to the PND (*P* > 0.01). Yellow boxes highlight the cancers with one of the two genes repositioned. Orange box highlights the cancer in with the two genes reposition, but in different directions.
