## Supplementary material for "Assessment of the Utility of Gene Positioning Biomarkers in the Stratification of Prostate Cancers": Sup Fig 1

### Supplementary Figure 1

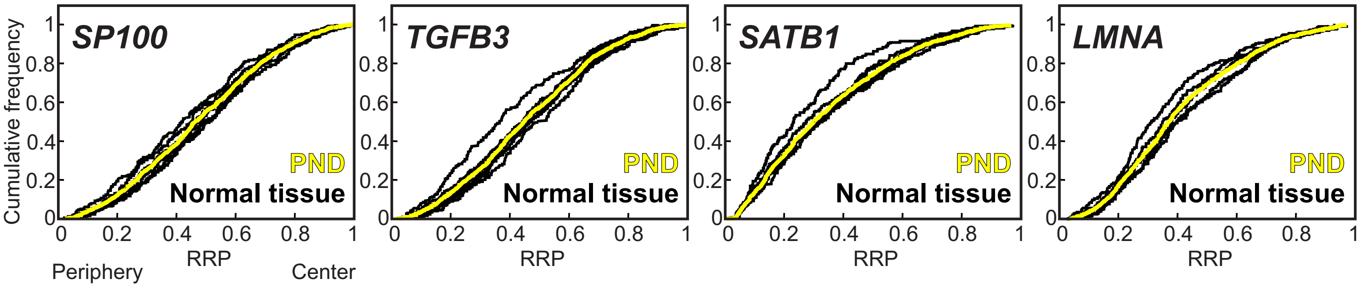


**Supplementary Figure 1. Pooled normal distribution.** Cumulative RRDs for individual normal prostate tissue (black) and the pooled normal distribution (PND; yellow), generated from combining all normal nuclei for a given gene into a single dataset.
