## Supplementary material for "Assessment of the Utility of Gene Positioning Biomarkers in the Stratification of Prostate Cancers": Sup Fig 2

### Supplementary Figure 2

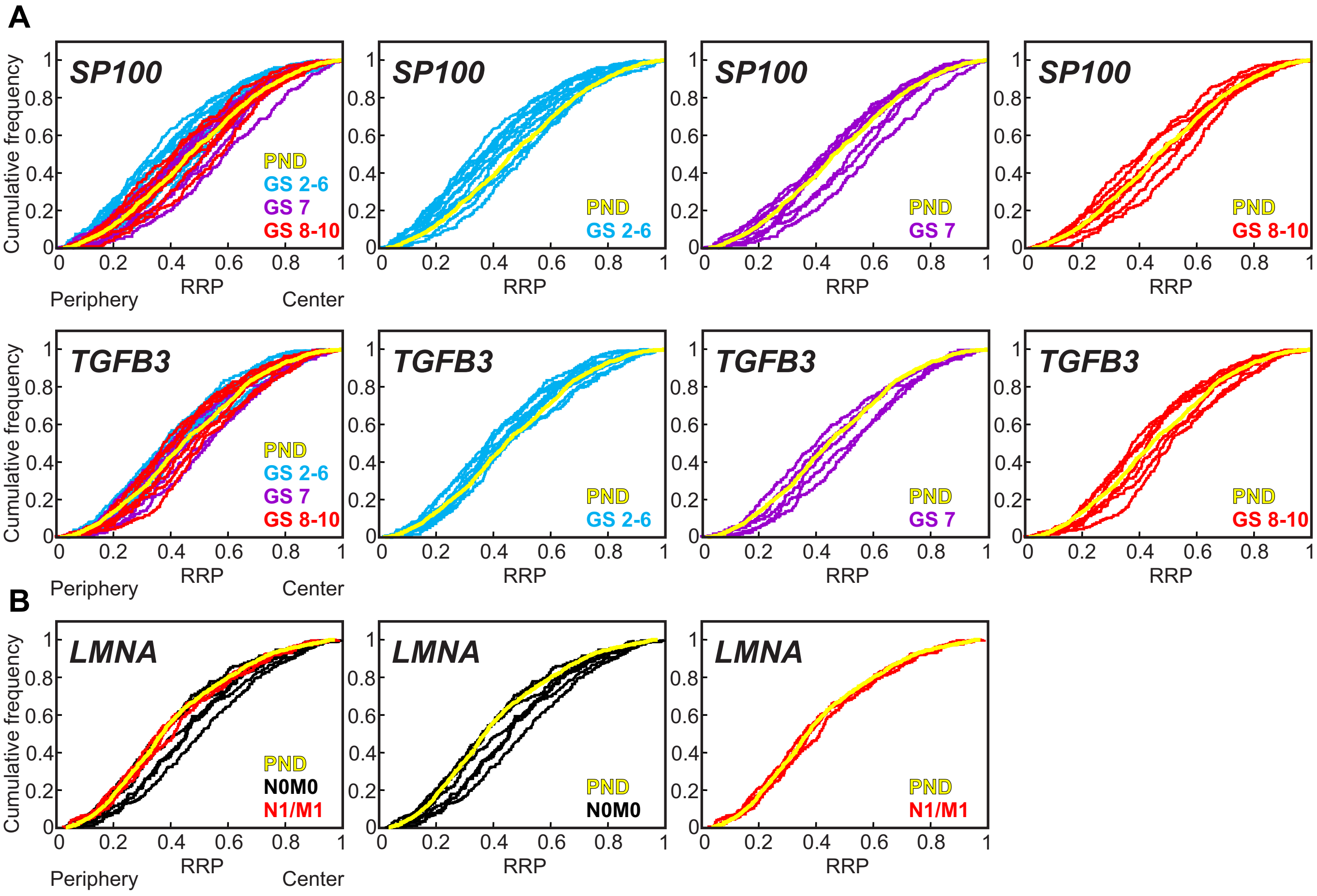


**Supplementary Figure 2. Sub-type specific spatial gene repositioning in cancer. (A)** Cumulative RRDs for *SP100* and *TGFB3* in prostate cancer tissues are color coded according to Gleason score: Gleason score 2-6 (blue), Gleason score 7 (purple) and Gleason score 8-10 (red). **(B)** Cumulative RRDs for *LMNA* in prostate cancer tissues are color coded according to metastatic status: Non-metastatic (N0M0) cancers (black), metastatic (N1/M1) cancers (red). In all graphs the pooled normal distribution (yellow) is included for comparison. GS, Gleason score; RRP, relative radial position.
